# Bacterial Diversity and Chemical Ecology of Natural Product-Producing Bacteria from Great Salt Lake Sediment

**DOI:** 10.1101/2023.11.07.565188

**Authors:** Elijah R. Bring Horvath, William J. Brazelton, Min Cheol Kim, Reiko Cullum, Matthew A. Mulvey, William Fenical, Jaclyn M. Winter

## Abstract

Great Salt Lake (GSL), located northwest of Salt Lake City, UT, is the largest terminal lake in the United States. While the average salinity of seawater is ∼3.3%, the salinity in GSL ranges between 5-28%. In addition to being a hypersaline environment, GSL also contains toxic concentrations of heavy metals, such as arsenic, mercury, and lead. The extreme environment of GSL makes it an intriguing subject of study, both for its unique microbiome and its potential to harbor novel natural product-producing bacteria, which could be used as resources for the discovery of biologically active compounds. Though work has been done to survey and catalogue bacteria found in GSL, the Lake’s microbiome is largely unexplored, and little-to-no work has been done to characterize the natural product potential of GSL microbes. Here, we investigate the bacterial diversity of two important regions within GSL, describe the first genomic characterization of Actinomycetota isolated from GSL sediment, including the identification of a new *Saccharomonospora* species, and provide the first survey of the natural product potential of GSL bacteria.

## Introduction

Great Salt Lake (GSL), located in northern Utah, USA, is the largest saltwater body in the United States. Its area spans five counties, including Weber, Box Elder, Salt Lake, Tooele, and Davis, and covers approximately 4,400 km^2^, though its area is rapidly shrinking [1, 2, 3]. Of ecological importance, GSL is a major bird nesting spot, hosting millions of migratory birds each year, which are a necessary component of the complex GSL ecosystem [3]. GSL is additionally a terminal lake, meaning that water flowing into it only leaves by evaporation, and consequently minerals, ions, and salts that enter the lake are retained and concentrated. A railroad causeway constructed in the 1950s physically divides GSL into two main regions: the North and South Arms. Because of this, salinity often reaches ≥ 27% in the North Arm, which receives little to no fresh water inflow, and it can range from 5% to greater than 15% in the South Arm [4]. A broader microbial diversity is observed in the South Arm compared to the North Arm, and this is primarily attributed to the differences in salinity [5, 6]. Aside from being a hypersaline environment, GSL also exhibits elevated levels of heavy metals and metalloids, including mercury, arsenic, cadmium, and lead [7]. Due to the extreme environments of GSL, microorganisms surviving and thriving in the North and South Arms have been described as polyextremophiles, as they have adapted to not only toxic levels of heavy metals and extreme osmotic stress, but also to extreme seasonal temperature fluctuations [3, 5]. To thrive in these harsh environments, GSL microorganisms may evolve and employ different methods for survival, such as heavy metal efflux, degradation of toxic compounds [8], and production of specialized small molecules. Of particular interest to us is the natural product potential of GSL bacteria.

Natural products are small molecules produced in nature and represent some of the most important pharmaceutical agents used in human health care [9, 10, 11]. This especially holds true with anti-infective agents, as many of the current clinically significant antibiotics are natural products or derivatives thereof. However, antibiotic resistance is on the rise and has been recognized by the World Health Organization as a leading global health issue [12, 13], which needs to be urgently addressed. According to the Center for Disease Control, 2.8 million Americans will acquire drug-resistant bacterial infections resulting in 35,000 deaths each year [14, 15], and the COVID-19 pandemic resulted in a drastic increase in antibiotic resistance-associated infections and deaths [16]. Antibiotic discovery over the last few decades has relied heavily on chemically modifying the scaffolds of known antibiotic agents. Therefore, the identification of bioactive natural products possessing new chemical scaffolds is a promising approach for the discovery of antibiotics with novel modes of action. Unique environments can influence the chemical diversity of natural products, and microorganisms isolated from extreme environments serve as ideal resources for drug discovery efforts. Furthermore, because GSL is a terminal lake and serves as the endpoint of wastewater treatment runoff from a major metropolitan center, we expect pathogenic microbes residing there to be a reservoir of antibiotic resistance genes. Therefore, native GSL microbes may be producing novel antibiotic agents to compete with these pathogens, which could make them an ideal resource for the identification of new natural products with novel scaffolds and new mechanisms of action.

To prioritize strains for downstream fermentation studies and streamline the labor-intensive natural product discovery pipeline when working with large numbers of bacterial isolates, robust bioinformatic and genomic tools can complement bioactivity-guided isolation efforts [17, 18, 19]. In the producing organism, the genes encoding the enzymatic machinery used to assemble small molecules are typically clustered together within a chromosome or on extra-chromosomal genetic elements. By correlating genetic information to protein function, chemical logic can be used to connect a natural product to its respective biosynthetic gene cluster [20]. However, little information on bacterial populations and even less on bacterial genomes has been reported from GSL [3, 5, 21], with most research focusing on planktonic communities isolated from water samples [22, 23, 24, 25, 26, 27, 28] rather than sediment-derived microbial populations [6]. Although previous studies have explored the microbial ecology of GSL using culture-independent methods [6, 26], no genomic studies have been published [21, 27, 28]. Thus, it is imperative that we further characterize this unique and understudied ecosystem, with a focus on expanding our knowledge of what sediment-dwelling microbes are present using both culture-dependent and independent methods. Just as importantly, we must also assess GSL bacteria for their potential to produce novel natural products. Due to severe drought-related shrinkage of the Lake, it is imperative we perform these studies before this invaluable resource is gone [1, 3]. Here, we investigate the microbial diversity in the South Arm of GSL, as well as characterize the natural product potential of the first Actinomycetota strains isolated from GSL sediment.

## Results

### Microbial Diversity of GSL Sediments in the South Arm

We collected eight sediment samples from two different geographic regions within the South Arm of GSL – Black Rock Beach (BRB) and the Marina, which are only ca. 200 yards apart from each other. From each region, four sediment samples were collected and assessed for the presence of bacteria using 16S rRNA gene amplicon sequencing, which resulted in the identification of 748,251 amplicon sequencing variants (ASVs) (BRB = 441,628 and Marina = 306,623). Though we expected to see little variation in bacterial community composition between our two geographically close collection sites, we found that the regions did exhibit significant differences in their taxonomic composition and structure (Figure 1; Bray-Curtis, p < 0.05, [PERMANOVA]). Upon phylogenomic analysis, we found that the ASVs comprised 53 phyla and 421 genera (Table 1, Table S1). We identified Pseudomonadota (49% of total ASVs) and Bacteroidota (22% of total ASVs) as the most abundant phyla, and Gammaproteobacteria (34% of total ASVs) and Bacteroidia (17% of total ASVs) as the most abundant classes (Table 1).

**Table 1.**
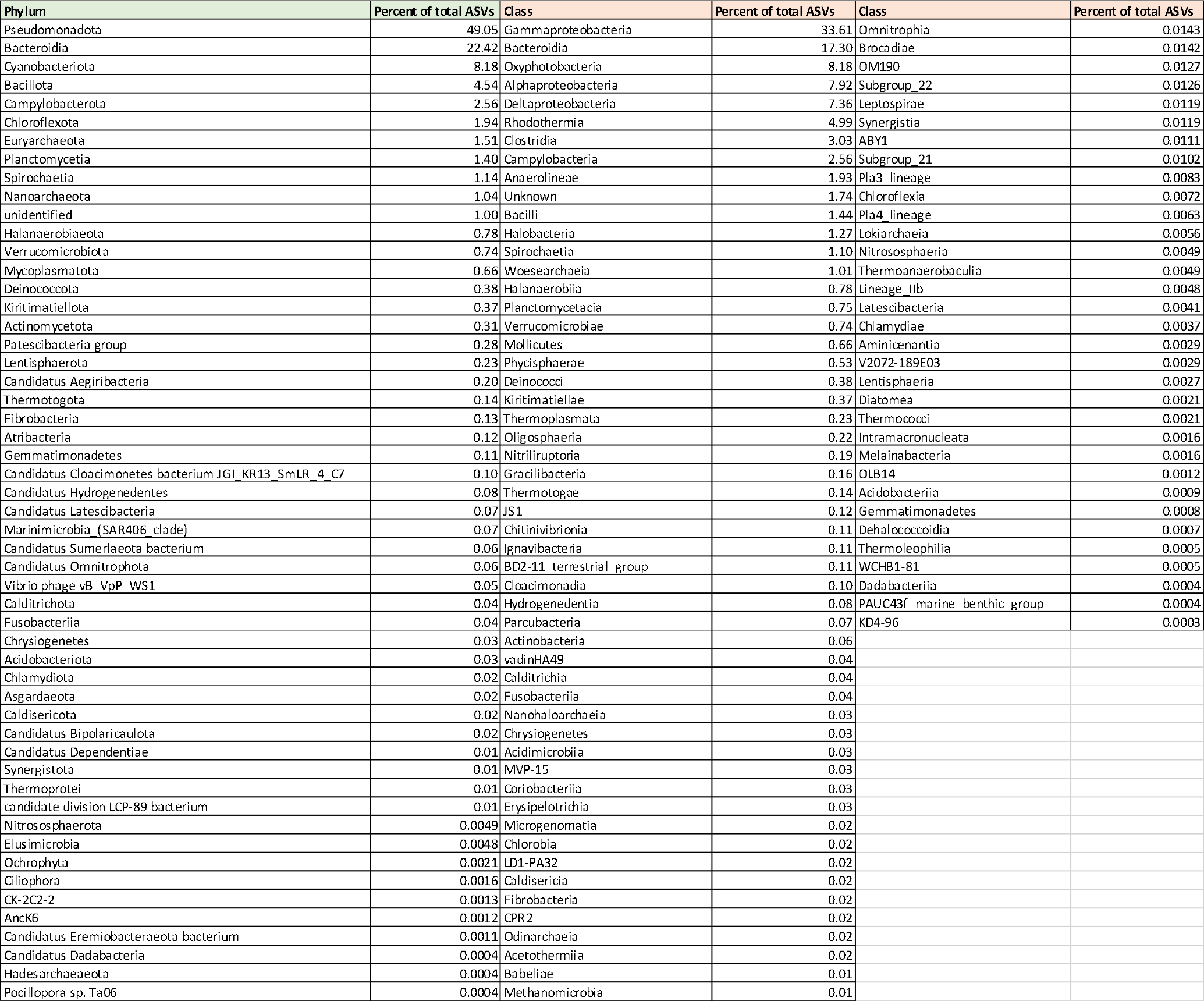
Diversity and amplicon sequence variant (ASV) abundance of GSL microbes.

**Figure 1.**
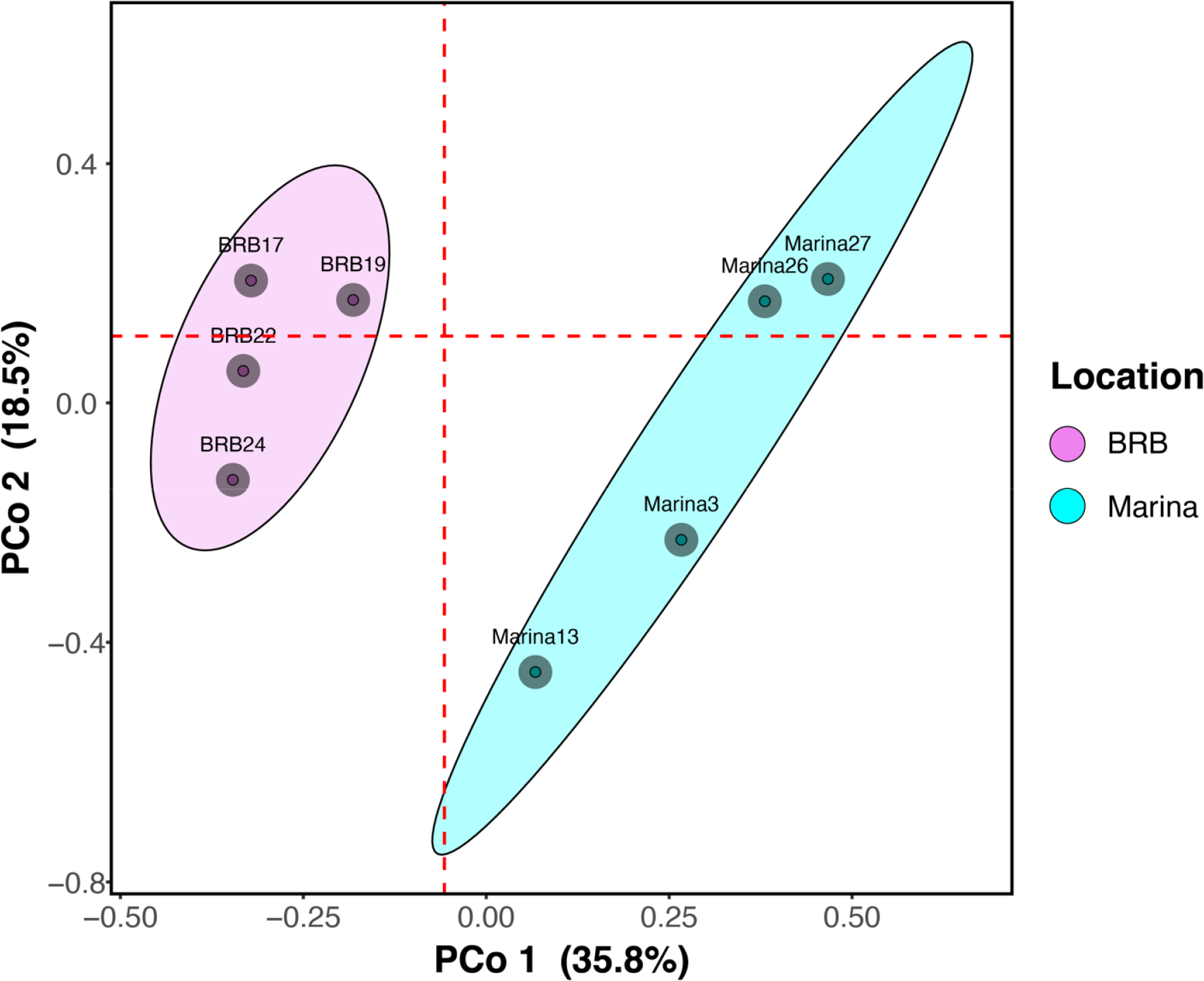
Bacterial community composition of sediment samples from Black Rock Beach and the Marina regions, as measured by 16S rRNA gene amplicon sequencing. The principal coordinate analysis (PCoA) plot illustrates overall community similarities and differences between four BRB and four Marina collection sites in the South Arm of GSL. Dashed red lines represent medians, and ellipses represent 75% confidence intervals around the samples from each region.

Upon closer inspection of our datasets, we found that the relative abundances of ASVs associated with specific organisms were more abundant in one of the two regions, either more abundant in BRB sediment or more abundant in Marina sediment. For example, ASVs associated with several Planctomycetota genera were found at significantly higher levels in the Marina sediment compared to BRB sediment (*Pirellula, p* = 3.68e-13, *p* = 2.9e-4; *Rubripirellula, p* = 1.25e-14 [Wald test]) (Figure 2). In total, ASVs comprising 24 distinct phyla and 60 genera exhibited significantly different abundance between our two collection regions (Figures 2 and S2). Interestingly, we observed differential abundances of ASVs representing two *Sulfurimonas* spp., with one found at much higher abundance in our BRB sample (*p* = 5.16e-11) and the other from the Marina sediment (*p* = 9.22e-11). Actinomycetota, which are common natural product producers, were found in both regions (0.31% of total ASVs), and an unknown *Saccharomonospora* species was found at a higher abundance in the Marina sediment compared to BRB (*p* = 5.24e-12) (Figure 2, Figure S1).

**Figure 2.**
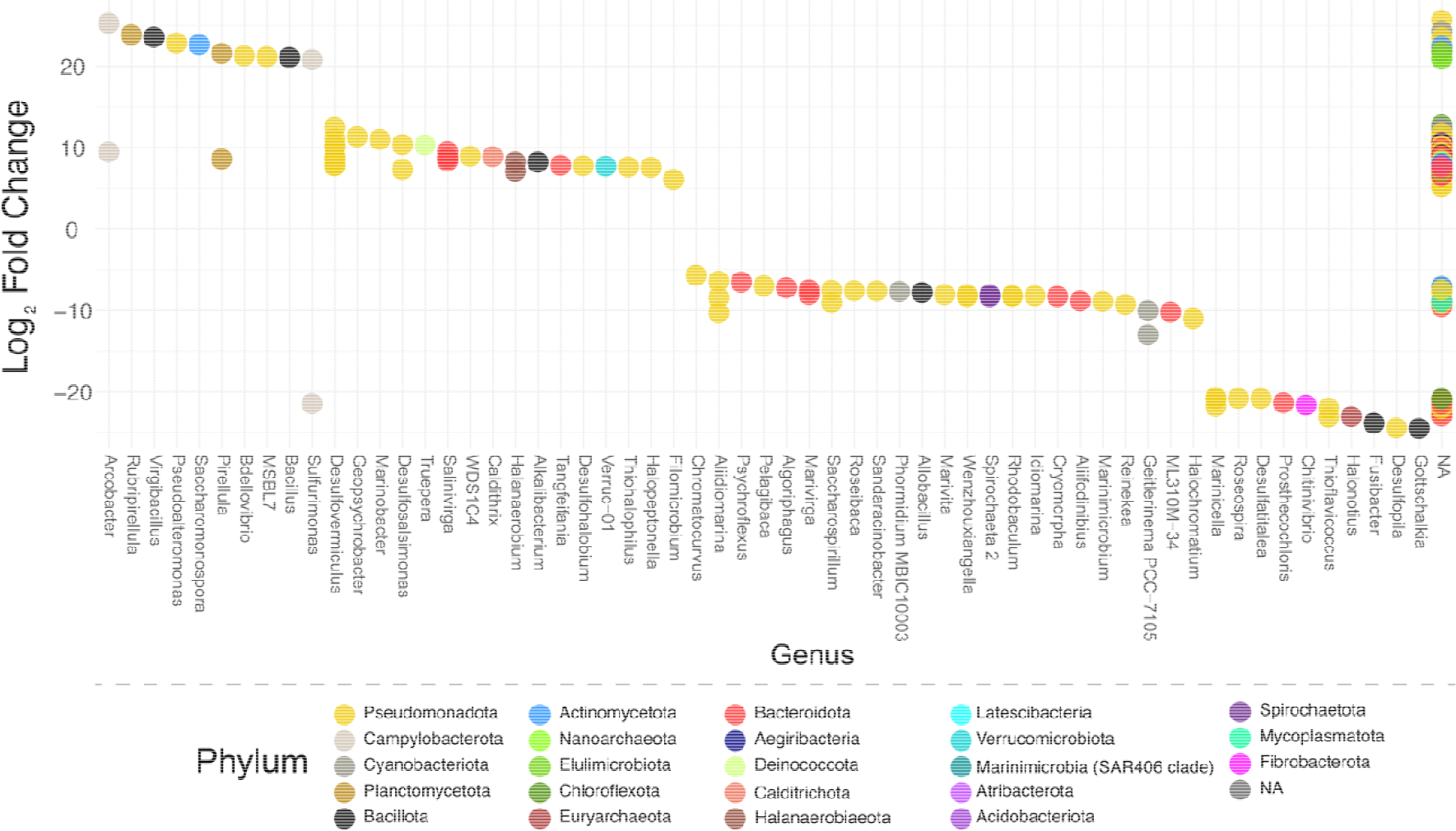
DESeq2 plot showing log_2_ fold change of phyla on the y-axis and genera on the x-axis illustrating differential abundance of microorganisms between Black Rock Beach and the Marina based on amplicon sequence variants (ASV). A positive log_2_ fold-change indicates a significantly higher abundance at the Marina, while a negative log_2_ fold-change indicates significantly higher abundance at Black Rock Beach. Analysis was generated using an adjusted p-value cutoff of < 0.01. NA/NA values represent unknown phylum/unknown genus, and potentially represent uncharacterized bacteria.

### Taxonomic Classification of GSL Actinomycetota Strains

Using a culture-dependent approach, we isolated and sequenced two Actinomycetota strains, *Saccharomonospora* sp. GSL17-019 and *Streptomyces* sp. GSL17-113 from sediment collected at the Marina. To determine the taxonomic position of *Saccharomonospora* sp. GSL17-019 and *Streptomyces* sp. GSL17-113, we conducted whole-genome phylogenomic analyses. Average nucleotide identity (ANI) and digital DNA-DNA hybridization (dDDH) analyses were conducted on each genome (Table S2) [29, 30, 31], and maximum-likelihood phylogenetic trees were assembled based on multi-locus sequence analysis [31] (Figure S3 and S4). Interestingly, while *Streptomyces* sp. GSL17-113 was closely related to Streptomyces albus, *Saccharomonospora* sp. GSL17-019 did not exhibit significant relatedness with any known *Saccharomonospora* species, with its closest related type-strain ranging from 51.8% to 57.2% identity (Table S2). As this dDDH value is well below the species threshold of 70%, we posit that strain GSL17-019 represents a new *Saccharomonospora* species that could be specific to GSL. In addition to whole-genome sequencing of *Saccharomonospora* sp. GSL17-019 and *Streptomyces* sp. GSL17-113, we also sequenced the metagenome of a mixed-population culture comprised of Streptomyces and Bacillus spp., termed GSL17-111M from Marina sediment.

### Natural Product Potential of GSL Bacteria

We investigated the potential for specialized metabolite production by our sequenced isolates through the identification and annotation of natural product biosynthetic genes clusters (BGCs) encoded in each genome (Figure 3, Tables S3-S5). We identified 20 putative BGCs in *Saccharomonospora* sp. GSL17-019 (Table S3) and 27 putative BGCs in *Streptomyces* sp. GSL17-113 (Table S4), representing 19 different BGC classes. These classes included more common polyketide synthase and non-ribosomal peptide synthetase-containing clusters, as well as less common ribosomally synthesized and post-translationally modified peptide clusters predicted to produce lassopeptide [32] and ranthipeptide [32, 33] natural products (Figure 3A). From our mixed GSL population, GSL17-111M, we identified more than 80 BGCs (Figure 3B, Table S5). Of the combined BGCs identified, only nine strongly correlated to characterized clusters in public databases that have been associated with specific natural products (Figure 4). To better assess the natural product potential of our GSL isolates, *Streptomyces* sp. GSL17-113 was subjected to small-scale cultivation studies. From the culture extract, we identified tambjamine BE-18591 (Figure 5B, S5, S6, and Table S6). Tambjamine BE-18591 was originally reported from *Streptomyces* sp. BA18591, a plant-derived isolate collected in Japan [34, 35]. The tambjamines possess antimicrobial and cytotoxic activities [34, 36], and tambjamine BE-18591 has been reported to possess antimicrobial activity against both fungi and bacteria, including *Candida albicans, Malassezia furfur, Escherichia coli*, and *Staphylococcus aureus* [34, 36]. Further, tambjamine BE-18591 displays broad antitumor activity against leukemia, melanoma, colorectal, and glioblastoma cell lines [36], as well as inhibition of immunoproliferation and gastritis in rabbits [37]. Our genomic analysis also confirmed the presence of the tambjamine BE-18591 BGC in *Streptomyces* sp. GSL17-113 (Figure 5A and S2, and Tables S6 and S7), emphasizing the importance and utility of genome mining when prioritizing strains for downstream fermentation studies and aiding in the dereplication process. Though our initial fermentation study resulted in the isolation of a known compound, more than 90% of the identified gene clusters (> 120 BGCs) annotated in our three datasets had weak-to-no similarity with characterized BGCs, indicating that the corresponding biosynthetic machinery could be synthesizing new natural product scaffolds.

**Figure 3.**
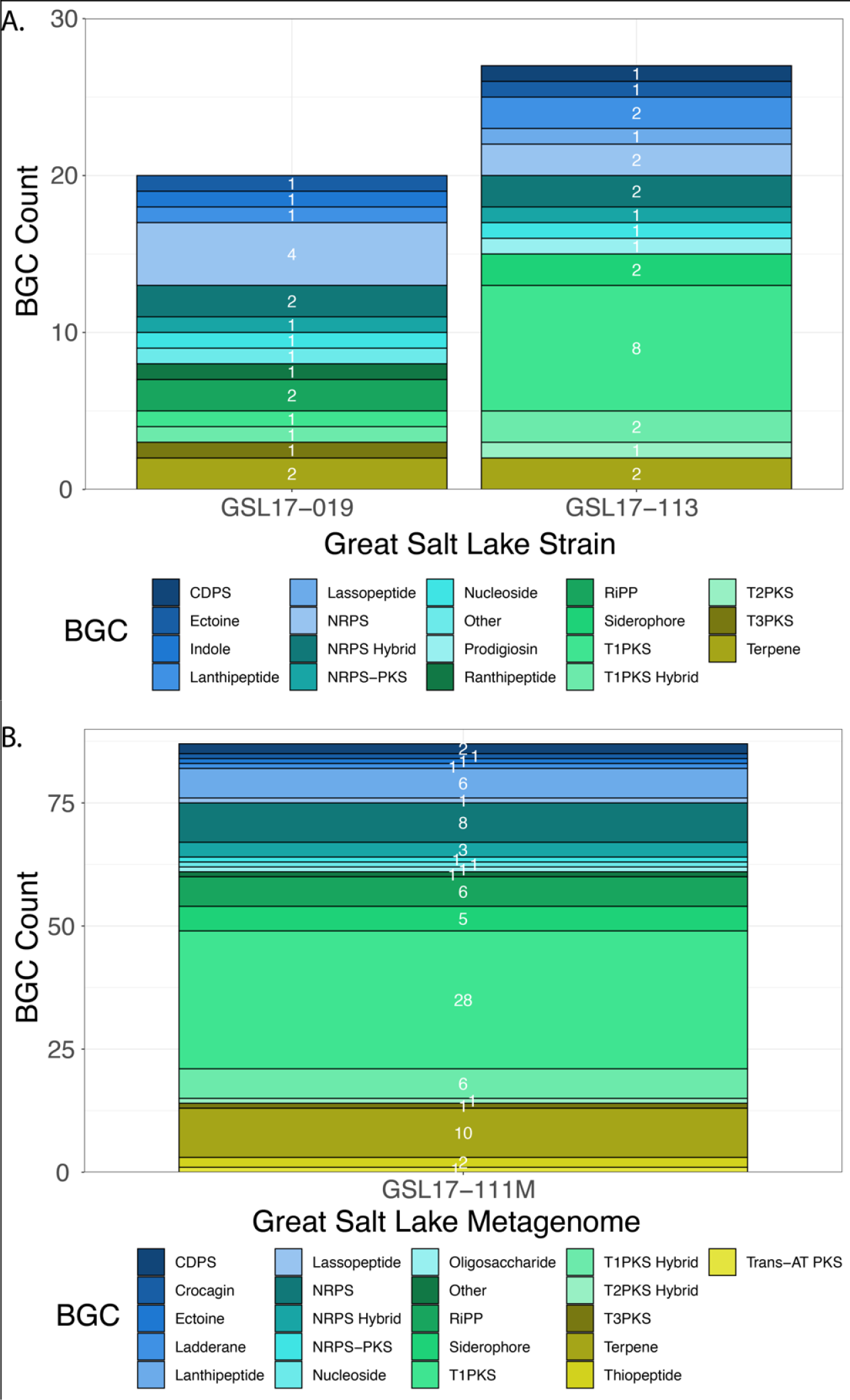
Natural product potential of GSL bacteria. A-B. Biosynthetic gene cluster (BGC) diversity of GSL bacterial strains. **A**. BGCs identified in *Saccharomonospora* sp. GSL17-019 and *Streptomyces* sp. GSL17-113. **B**. BGCs identified in GSL population genome, GSL17-111M, comprised of *Streptomyces* and *Bacillus* spp. Abbreviations: cyclodipeptide synthase (CDPS), non-ribosomal peptide synthetase (NRPS), polyketide synthase (PKS), ribosomally synthesized and post-translationally modified peptide (RiPP), acyltransferase (AT), type I polyketide synthase (T1PKS), type II polyketide synthase (T2PKS), type III polyketide synthase (T3PKS).

**Figure 4.**
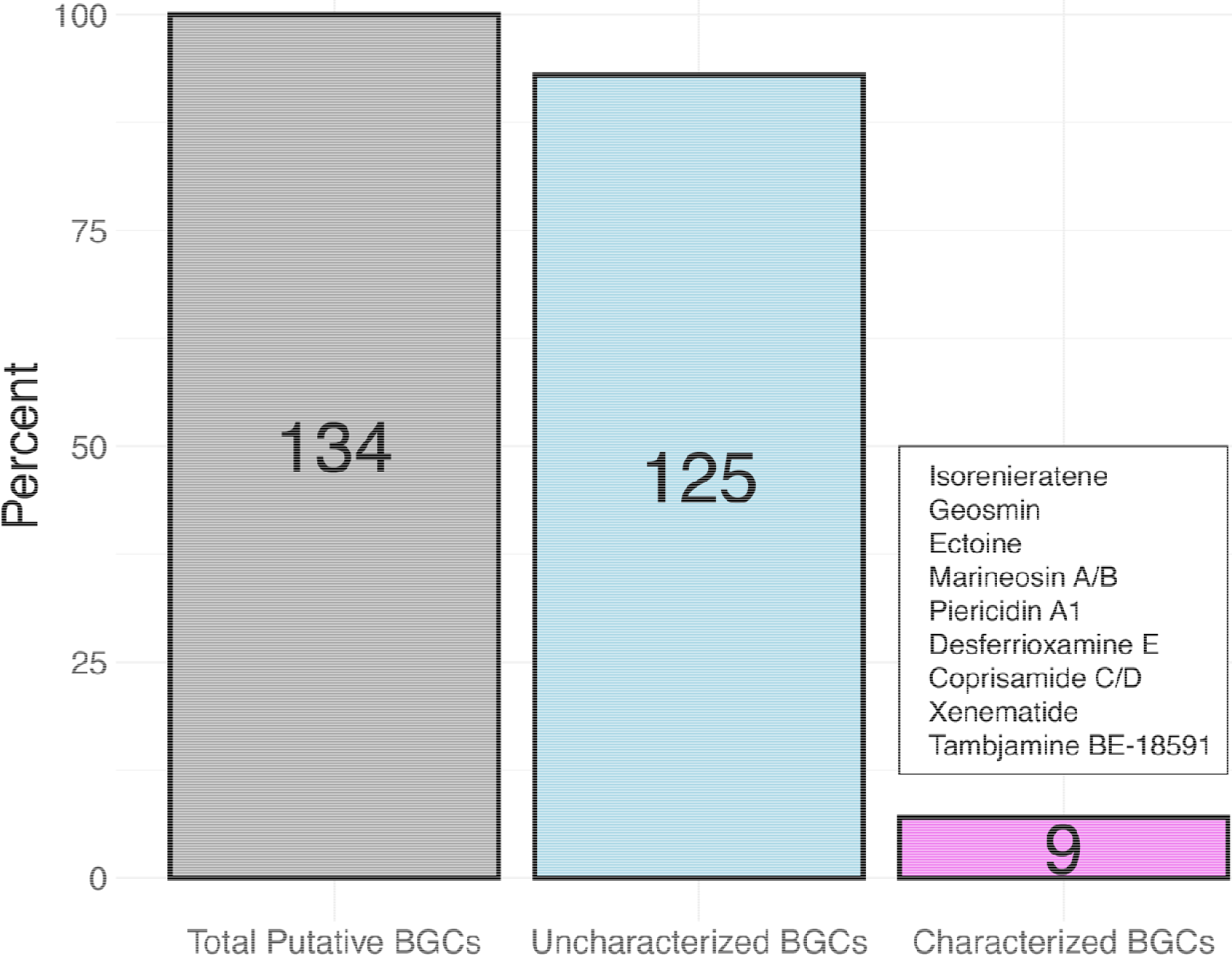
Comparison of all putative biosynthetic gene clusters (BGCs) identified in *Saccharomonospora* sp. GSL17-019, *Streptomyces* sp. GSL17-113, and the GSL population genome, GSL17-111M. A total of 134 BGCs were identified. Of those, nine were strongly associated with characterized BGCs deposited in MIBig (≥ 85% predicted similarity), while the remaining 125 BGCs were not strongly associated with known gene clusters. BGC-associated compounds were identified through gene cluster comparison using both antiSMASH and manual annotation and comparison of our BGCs to known BGCs.

**Figure 5.**
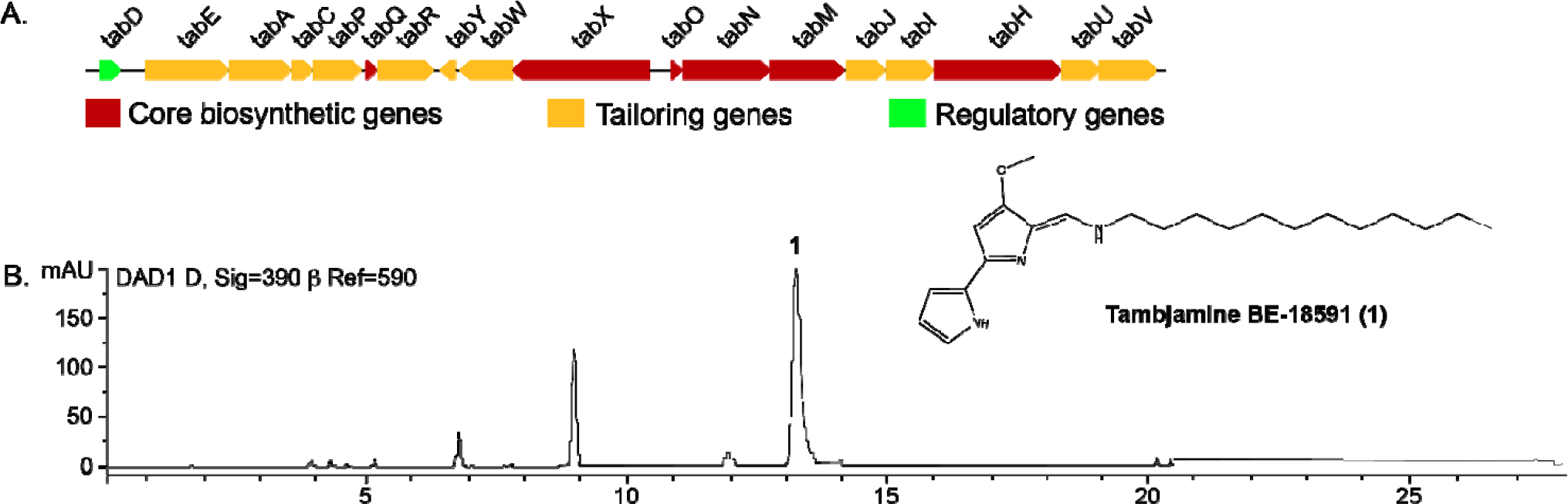
Identification of tambjamine BE-18591 in *Streptomyces* sp. GSL17-113. **A**. Organization of the tambjamine BE-18591 biosynthetic gene cluster. **B**. High performance liquid chromatography analysis (390 nm) of GSL17-113 and identification of tambjamine BE-18591 (**1**).

## Discussion

In this study, we have highlighted the diversity of GSL’s unique microbiome, including the genomic characterization of a potentially new *Saccharomonospora* species, and emphasized the natural product potential of GSL bacteria to produce bioactive compounds. Previously, we isolated natural products from a GSL organism, named the bonnevillamides. These compounds are linear heptapeptides containing a distinctive 3-(3,5-dichloro-4-hydroxyphenyl)-2-methoxypropenoic acid moiety, and bonnevillamide A harbors an additional unprecedented 4-methyl-azetidine-2-carboxylic acid methyl ester chemical motif [38, 39]. We also reported on the isolation and structure elucidation of the salinipeptins, ribosomally synthesized and post-translationally modified peptides containing rare D-amino acids, a highly functionalized N-terminus, and a C-terminal aminovinyl-cysteine residue [40]. Importantly, salinipeptin A displayed moderate activity against Group A *Streptococcus pyogenes*, as well as glioblastoma and colon cancer cell lines. Although tambjamine BE-18591 is a known natural product, its isolation from *Streptomyces* sp. GSL17-113 represents an additional natural product exhibiting antimicrobial and antitumor activities isolated from GSL. This further supports GSL as an underexplored resource for bioactive natural products and additional fermentation studies with both *Streptomyces* sp. GSL17-113 and *Saccharomonospora* sp. GSL17-019 are underway.

In parallel with the isolation and characterization of novel natural products, we intend to explore potential resistance mechanisms utilized by GSL microbes. As GSL is a terminal lake and serves as the endpoint for wastewater treatment runoff, we anticipate it to be a significant source of antibiotic-resistant pathogenic microbes. Indeed, the Gammaproteobacteria class within the Pseudomonadota phylum, encompassing well-characterized bacterial pathogens like *Escherichia coli, Klebsiella pneumoniae*, and *Pseudomonas aeruginosa* were the most abundant class of bacteria identified at both the Black Rock Beach and Marina regions, representing 34% of total ASVs (Table 1). Antibiotic resistance poses a monumental health challenge, and the environment can serve as a reservoir for antibiotic-resistant bacteria [41, 42, 43, 44, 45, 46]. Though it will require additional work to address this pressing concern, it is important that we thoroughly document patterns of resistance found in microbes within our local environments, as well as work to develop anti-infective agents to combat multidrug-resistant microorganisms. Furthermore, from an ecological standpoint, as GSL water levels fluctuate, it is important we continue to investigate the Lake’s microbiome and how changing water levels, drought, and inflow affect microbial diversity. From this study, we not only provide initial insight into the diversity of GSL sediment-derived bacteria, but also provide the first genomic characterization of Actinomycetota isolated from GSL. This is of particular interest to our group, as Actinomycetota are the dominant producers of therapeutic compounds essential for human health [44, 47, 48, 49].

Bioprospecting of GSL bacteria led to the identification of the bonnevillamides, salinipeptins, and in this study, the antitumor antibiotic compound, tambjamine BE-18591. Given the numerous unidentified biosynthetic gene clusters (Figure 4, Table S3, S4, and S5), it is evident that GSL represents a largely untapped resource of bioactive natural products. Though our primary interest is in characterizing known natural product producing microbes, the high number of unknown phylum/unknown genus values we observed in our sediment samples (Figures 2 and 3) suggests that GSL is home to many as-of-yet characterized bacterial organisms. This is emphasized by the discovery of the potentially new *Saccharomonospora* species GSL17-019. Taken together, our results give an overview of GSL’s large microbial and chemical diversity, as well as unexpected differences in microbial populations between geographically close regions and revealed GSL’s potential to harbor new species of natural product-producing bacteria.

## Materials and Methods

### Collection and Isolation of Bacteria Strains

Sediment samples were collected from the South Arm of GSL at four Black Rock Beach and four Marina sites during the summer of 2017. *Saccharomonospora* sp. GSL17-019 and *Streptomyces* sp. GSL17-113 were isolated from Marina sediment that was desiccated for 72 hours in a biological safety cabinet. Desiccated sediment, ∼0.5 g, was added to yeast-peptone-mannitol (YPM) agar plates (per liter: 4 g mannitol, 2 g yeast extract, 2 g peptone, 18 g agar, and 42 g Instant Ocean Aquarium Sea Salt Mixture, Spectrum Brands, USA). Plates were incubated at 30°C for up to 90 days, and bacterial colonies were subcultured on YPM media until pure isolates were obtained (with the exception of GSL17-111M). 16S rRNA gene sequencing was used to identify *Saccharomonospora* sp. GSL17-019, *Streptomyces* sp. GSL17-113, and the constituents of GSL17-111M using NCBI Blast/Blast+ [50] (GenBank accession for reference 16S rRNA sequence is CP082835.1).

### Environmental Sequencing of 16S rRNA Genes

DNA preparations of Great Salt Lake sediment were carried out using a FastDNA Spin Kit (MP Biomedicals, cat. no.: 116540-600). Sequencing of amplicons generated from 16S rRNA genes was performed at the Genomics Core Facility at Michigan State University on an Illumina MiSeq instrument using dual-indexed Illumina fusion primers targeting the V4 region of the 16S rRNA gene [51]. Amplicon sequence variants (ASVs) were inferred from 16S rRNA amplicon sequences with DADA2 v. 1.10.1 [52] after removal of primer sequences with cutadapt v. 1.15 [53]. Taxonomic classification of ASVs was performed with DADA2 using the SILVA reference alignment (SSURefv132) and taxonomy outline [54, 55]. The ordination plot was generated with PhyloSeq v. 1.26.1 [19] using PCoA ordination of Bray-Curtis dissimilarity values.

### Genome Sequencing, Genome Mining, and Phylogenomic Analyses of GSL Genomes

Construction of genomic libraries and sequencing was performed at the High-Throughput Genomics Center in the Huntsman Cancer Institute at the University of Utah. A PCR-free NEBNext Ultra II DNA library was generated and sequenced using NovaSeq S4 Reagent Kit v1.5 (150×150 bp and 2500 M read-pairs/lane). Adapter sequences and PhiX were removed from all reads with BBDuk, part of the BBtools suite, V35.85 [56]. Quality trimming was performed with seq-qc as previously described [57], and sequences were assembled using SPAdes [58]. Biosynthetic gene clusters were initially identified using antiSMASH [17], and clusters of interest were manually annotated and characterized using NCBI Blast/Blast+. Biosynthetic gene clusters (BGCs) identified in our GSL genomes that exhibited ≥ 85% similarity to characterized BGCs in the MIBiG [59] database were classified as ‘known’ in our analyses. Phylogenomic analyses, including average nucleotide identity (ANI) and digital DNA-DNA hybridization (dDDH) were conducted using autoMLST [31] and the Type Strain Genome Server (TYGS) [29, 30].

### Statistical Analysis and Data Visualization

NMDS plots and DESeq2 analyses were carried out using PhyloSeq v. 1.42.0 [18, 19] in RStudio v. 2023.3.0.386 [60]. DESeq2 v. 1.38.3 was used to compare amplicon sequence variant data from the four Black Rock Beach (BRB) and four Marina sampling sites and graph differential abbundance. During the DESeq2 analysis, each of the four collection sites from both BRB and the Marina were treated as replicates of the primary collection region to obtain a BRB vs Marina comparison. Significance was determined using a Wald test with a Benjamini and Hochberg adjusted p-value. Only amplicon sequence variants (ASVs) meeting the minimal cut-off of an adjusted p-value of p > 0.01 were included in the results. Bar plots were created in RStudio using ggplot2 v. 3.4.2 and ggbreak v. 0.1.1 [61]. PhyloSeq-generated taxonomy classifications were updated to reflect current literature using the NCBI taxonomy browser. Phylogenetic trees were constructed with autoMLST using IQ-TREE Ultrafast Bootstrap analysis of 1000 replicates. The autoMLST-generated phylogenetic tree was exported as a Newick file and used to generate final figures in R using ggtree v.3.8.0 [62].

## Data Availability

All relevant code and data are available at https://github.com/ERBringHorvath/GSL_Phyloseq. 16S amplicon sequences are available under BioProject accession PRJNA975952.

## Acknowledgements

This work was supported by a 3iii graduate research fellowship to ERBH, by the US National Institutes of Health (1R01AI155694) to WF and JMW, by the University of Utah Research Foundation to JMW, by the Margolis Foundation to WJB, MAM and JMW, and in part by the Department of Defense award (W81XWH-22-1-0800, SC210103) to WJB, JMW, and MAM. We thank Abby Scott, University of Utah, for assistance with processing GSL sediment and helpful discussions.

## References

1. Carlowicz M. Record low for Great Salt Lake. NASA Earth Observatory. 2021. https://earthobservatory.nasa.gov/images/148700/record-low-for-great-salt-lake. Accessed 31 Oct 2023.

2. Yeung P. Salt lake confronts a future without a lake. In: Bloomberg. 2022. https://www.bloomberg.com/news/features/2022-07-08/drought-leaves-salt-lake-city-with-a-looming-water-crisis?utm_source=website&utm_medium=share&utm_campaign=copy. Accessed 31 Oct 2023.

3. Baxter BK. Great Salt Lake microbiology: A historical perspective. Int Microbiol. 2018; 21:79–95.

4. Merck MF, Tarboton DG. The salinity of the Great Salt Lake and its deep brine layer. Water. 2023; 15:1488.

5. Baxter BK, Zalar P. The extremophiles of Great Salt Lake: Complex microbiology in a dynamic hypersaline ecosystem. In: Seckbach, J, Rampelotto, P, (eds) Astrobiology exploring life on earth and beyond, model eccosystems in extreme environments. Academic Press, 2019. pp. 57–99.

6. Lindsay MR, Anderson C, Fox N, Scofield G, Allen J, Anderson E, et al. Microbialite response to an anthropogenic salinity gradient in Great Salt Lake, Utah. Geobiology. 2017; 15:131–45.

7. Ekrami J, Nemati Mansour S, Mosaferi M, Yamini Y. Environmental impact assessment of salt harvesting from the salt lakes. J Environ Health Sci Eng. 2021; 19:365–77.

8. Shuaib M, Azam N, Bahadur S, Romman M, Yu Q, Xuexiu C. Variation and succession of microbial communities under the conditions of persistent heavy metal and their survival mechanism. Microb Pathog. 2021; 150:104713.

9. Newman DJ, Cragg GM. Natural products as sources of new drugs over the nearly four decades from 01/1981 to 09/2019. J Nat Prod. 2020; 83:770–803.

10. Dias DA, Urban S, Roessner U. A historical overview of natural products in drug discovery. Metabolites. 2012; 2:303–36.

11. Hutchings MI, Truman AW, Wilkinson B. Antibiotics: Past, present and future. Curr Opin Microbiol. 2019; 51:72–80.

12. Darby EM, Trampari E, Siasat P, Gaya MS, Alav I, Webber MA, et al. Molecular mechanisms of antibiotic resistance revisited. Nat Rev Microbiol. 2022; 21:280–95.

13. Langford BJ, Soucy J-PR, Leung V, So M, Kwan ATH, Portnoff JS, et al. Antibiotic resistance associated with the covid-19 pandemic: A systematic review and meta-analysis. Clin Microbiol Infect. 2023; 29:302–9.

14. CDC. Antibiotic resistance threats in the united states. US Department of Health and Human Services, CDC. 2019.

15. Centers for Disease Control and Prevention UDoHaHS. National infection & death estimates for antimicrobial resistance: Centers for Disease Control and Prevention, National Center for Emerging and Zoonotic Infectious Diseases (NCEZID), Division of Healthcare Quality Promotion (DHQP); 2021. https://www.cdc.gov/drugresistance/national-estimates.html#print.

16. CDC. Covid-19: U.S. Impact on antimicrobial resistance, special report 2022. US Department of Health and Human Services, CDC. 2022.

17. Blin K, Shaw S, Steinke K, Villebro R, Ziemert N, Lee SY, et al. Antismash 5.0: Updates to the secondary metabolite genome mining pipeline. Nucleic Acids Res. 2019; 47:W81–W7.

18. Love MI, Huber W, Anders S. Moderated estimation of fold change and dispersion for rna-seq data with deseq2. Genome Biol. 2014; 15:550.

19. McMurdie PJ, Holmes S. Phyloseq: An R package for reproducible interactive analysis and graphics of microbiome census data. PLoS One. 2013; 8:e61217.

20. Biermann F, and Helfrich, E. J. N. Hidden treasures: Microbial natural product biosynthesis off the beaten path. mSystems. 2021; 6:e00846–21.

21. Almeida-Dalmet S, Sikaroodi M, Gillevet PM, Litchfield CD, Baxter BK. Temporal study of the microbial diversity of the north arm of Great Salt Lake, Utah, U.S. Microorganisms. 2015; 3:310–26.

22. Daniels LL. On the flora of Great Salt Lake. Am Nat. 1917; 51:499–506.

23. Packard AS. The sea-weeds of salt lake. Am Nat. 1879; 13:701–03.

24. Vorhies CT. Notes of the fauna of the Great Salt Lake. Am Nat. 1917; 51:494–99.

25. ZoBell CE. Direct microscopic evidence of an autochthonous bacterial flora in Great Salt Lake. Ecology. 1937; 18:453–58.

26. Meuser JE, Baxter BK, Spear JR, Peters JW, Posewitz MC, Boyd ES. Contrasting patterns of community assembly in the stratified water column of great salt lake, utah. Microb Ecol. 2013; 66:268–80.

27. Parnell JJ, Rompato G, Latta LCt, Pfrender ME, Van Nostrand JD, He Z, et al. Functional biogeography as evidence of gene transfer in hypersaline microbial communities. PLoS One. 2010; 5:e12919.

28. Tazi L, Breakwell DP, Harker AR, Crandall KA. Life in extreme environments: Microbial diversity in Great Salt Lake, Utah. Extremophiles. 2014; 18:525–35.

29. Jan P Meier-Kolthoff AFA, Hans-Peter Klenk, Markus Göker. Genome sequence-based species delimitation with confidence intervals and imporoved distance functions. BMC Bioinform. 2013; 14:60.

30. Meier-Kolthoff JP, Goker M. Tygs is an automated high-throughput platform for state-of-the-art genome-based taxonomy. Nat Commun. 2019; 10:2182.

31. Alanjary M, Steinke K, Ziemert N. Automlst: An automated web server for generating multi-locus species trees highlighting natural product potential. Nucleic Acids Res. 2019; 47:W276–W82.

32. Montalban-Lopez M, Scott TA, Ramesh S, Rahman IR, van Heel AJ, Viel JH, et al. New developments in ripp discovery, enzymology and engineering. Nat Prod Rep. 2021; 38:130–239.

33. Chen Y, Wang J, Li G, Yang Y, Ding W. Current advancements in sactipeptide natural products. Front Chem. 2021; 9:595991.

34. Kojiri K NS, Suzuki H, Okura A, Suda H. A new antitumor substance, be-18591, produced by a streptomycete. I. Fermentation, isolation, physicochemical and biological properties. J Antibiot. 1993; 12:1799–803.

35. Nakajima S KK, and Suda H. A new antitumor substance, be-18591, produced by a streptomycete. II. Structure determination. J Antibiot. 1993; 46:1894–6.

36. Pinkerton DM, Banwell MG, Garson MJ, Kumar N, de Moraes MO, Cavalcanti BC, et al. Antimicrobial and cytotoxic activities of synthetically derived tambjamines c and e–j, be-18591, and a related alkaloid from the marine bacterium Pseudoalteromonas tunicata. Chem Biodivers. 2010; 7:1311–24.

37. Tanigaki K, Sato T, Tanaka Y, Ochi T, Nishikawa A, Nagai K, et al. Be-18591 as a new h+/cl− symport ionophore that inhibits immunoproliferation and gastritis. FEBS Lett. 2002; 524:37–42.

38. Wu G, Nielson JR, Peterson RT, Winter JM. Bonnevillamides, linear heptapeptides isolated from a Great Salt Lake-derived *Streptomyces* sp. Mar Drugs. 2017; 15:195.

39. Shin YH, Ban YH, Shin J, Park IW, Yoon S, Ko K, et al. Azetidine-bearing non-ribosomal peptides, bonnevillamides d and e, isolated from a carrion beetle-associated actinomycete. J Org Chem. 2021; 86:11149–59.

40. Shang Z, Winter JM, Kauffman CA, Yang I, Fenical W. Salinipeptins: Integrated genomic and chemical approaches reveal unusual D-amino acid-containing ribosomally synthesized and post-translationally modified peptides (ripps) from a Great Salt Lake *Streptomyces* sp. ACS Chem Biol. 2019; 14:415–25.

41. Bleichenbacher S, Stevens MJA, Zurfluh K, Perreten V, Endimiani A, Stephan R, et al. Environmental dissemination of carbapenemase-producing enterobacteriaceae in rivers in Switzerland. Environ Pollut. 2020; 265:115081.

42. Antunes P, Machado J, Sousa JC, Peixe L. Dissemination of sulfonamide resistance genes (sul1, sul2, and sul3) in Portuguese Salmonella enterica strains and relation with integrons. Antimicrob Agents Chemother. 2005; 49:836–9.

43. Centers for Disease Control and Prevention UDoHaHS. Antibiotic resistance threats in the united states. 2019.

44. D’Costa VM, King CE, Kalan L, Morar M, Sung WW, Schwarz C, et al. Antibiotic resistance is ancient. Nature. 2011; 477:457–61.

45. de Vries LE, Valles Y, Agerso Y, Vaishampayan PA, Garcia-Montaner A, Kuehl JV, et al. The gut as reservoir of antibiotic resistance: Microbial diversity of tetracycline resistance in mother and infant. PLoS One. 2011; 6:e21644.

46. Heather K. Allen JD, Helena Huimi Wang, Karen A. Cloud-Hansen, Julian Davies and Jo Handelsman. Call of the wild: Antibiotic resistance genes in natural environments. Nat Rev. 2010; 8:251–9.

47. Wainwright M. Streptomycin: Discovery and resultant controversy. Hist Phil Life Sci. 1991; 13:97–124. 48.

48. Lenz KD, Klosterman KE, Mukundan H, Kubicek-Sutherland JZ. Macrolides: From toxins to therapeutics. Toxins. 2021; 13:347.

49. Maiese WM, Lechevalier MP, Lechevalier HA, Korshalla J, Kuck N, Fantini A, Wildey MJ, Thomas J, Greenstein M. Calicheamicins, a novel familiy of antitumor antibiotics: Taxonomy, fermentation and biological properties. J Antibiot. 1989; 42:558–63.

50. Camacho C, Coulouris G, Avagyan V, Ma N, Papadopoulos J, Bealer K, et al. Blast+: Architecture and applications. BMC Bioinformatics. 2009; 10:421.

51. Kozich JJ, Westcott SL, Baxter NT, Highlander SK, Schloss PD. Development of a dual-index sequencing strategy and curation pipeline for analyzing amplicon sequence data on the miseq illumina sequencing platform. Appl Environ Microbiol. 2013; 79:5112–20.

52. Callahan BJ, McMurdie PJ, Rosen MJ, Han AW, Johnson AJ, Holmes SP. Dada2: High-resolution sample inference from illumina amplicon data. Nat Methods. 2016; 13:581–3.

53. Martin M. Cutadapt removes adapter sequences from high-throughput sequencing reads. EMBnet. 2011; 17.

54. Quast C, Pruesse E, Yilmaz P, Gerken J, Schweer T, Yarza P, et al. The silva ribosomal rna gene database project: Improved data processing and web-based tools. Nucleic Acids Res. 2013; 41:D590–6.

55. Yilmaz P, Parfrey LW, Yarza P, Gerken J, Pruesse E, Quast C, et al. The silva and “all-species living tree project (ltp)” taxonomic frameworks. Nucleic Acids Res. 2014; 42:D643–8.

56. Bushnell B, Rood J, Singer E. Bbmerge - accurate paired shotgun read merging via overlap. PLoS One. 2017; 12:e0185056.

57. Thornton CN, Tanner WD, VanDerslice JA, Brazelton WJ. Localized effect of treated wastewater effluent on the resistome of an urban watershed. Gigascience. 2020; 9:giaa125.

58. Bankevich A, Nurk S, Antipov D, Gurevich AA, Dvorkin M, Kulikov AS, et al. Spades: A new genome assembly algorithm and its applications to single-cell sequencing. J Comput Biol. 2012; 19:455–77.

59. Terlouw BR, Blin K, Navarro-Munoz JC, Avalon NE, Chevrette MG, Egbert S, et al. Mibig 3.0: A community-driven effort to annotate experimentally validated biosynthetic gene clusters. Nucleic Acids Res. 2023; 51:D603–D10.

60. Posit. Rstudio: Integrated development environment for R. Posit Softwar, PBC: Posit Softwar, PBC; 2023.

61. Xu S, Chen M, Feng T, Zhan L, Zhou L, Yu G. Use ggbreak to effectively utilize plotting space to deal with large datasets and outliers. Front Genet. 2021; 12:774846.

62. Yu G SD, Zhu H, Guan Y, Lam TT. Ggtree: An R package for visualization and annotation of phylogenetic trees with their covariates and other associated data. Methods Eco Evol. 2017; 8:28–36.

