## Supplemental Information for "Bacterial Diversity and Chemical Ecology of Natural Product-Producing Bacteria from Great Salt Lake Sediment"


**Author Information**

Elijah R. Bring Horvath,[1,2,3] William J. Brazelton,[2] Min Cheol Kim,[4] Reiko Cullum,[4] Matthew A. Mulvey,[2,5] William Fenical,[4,6,7] Jaclyn M. Winter[3,*]

[1]Department of Medicinal Chemistry, University of Utah, Salt Lake City, United States; [2]School of Biological Sciences, University of Utah, Salt Lake City, United States; [3]Department of Pharmacology and Toxicology, University of Utah, Salt Lake City, United States; [4]Center for Marine Biotechnology and Biomedicine, Scripps Institution of Oceanography, University of California at San Diego, San Diego, United States; [5]Henry Eyring Center for Cell and Genome Science, University of Utah, Salt Lake City, United States; [6]Skaggs School of Pharmacy and Pharmaceutical Sciences, University of California at San Diego, San Diego, United States; [7]Moores Comprehensive Cancer Center, University of California at San Diego, San Diego, United States.

### Table of Contents

**Supplementary Tables**
Table S1: DADA2 output and taxonomic classification of amplicon sequence variants **S3**
Table S2: ANI and dDDH results for *Saccharomonospora* sp. GSL17-019 and *Streptomyces* sp. GSL17-113 **S6**
Table S3: antiSMASH results of *Saccharomonospora* sp. GSL17-019 **S9**
Table S4: antiSMASH results of *Streptomyces* sp. GSL17-113 **S10**
Table S5: antiSMASH results of GSL population genome GSL17-111M **S11**
Table S6: $^1$H NMR assignment of tambjamine BE-18591 **S14**
Table S7: Putative tambjamine BE-18591 biosynthetic cluster in *Streptomyces* sp. GSL17-113 **S17**

**Supplementary Figures**
Figure S1: Differential abundance of ASVs associated with Actinomycetota and Pseudomonadota **S4**
Figure S2: DESeq2 analysis of differentially abundant ASVs **S5**
Figure S3: Maximum-likelihood phylogenetic tree of *Saccharomonospora* sp. GSL17-019 **S7**
Figure S4: Maximum-likelihood phylogenetic tree of *Streptomyces* sp. GSL17-113 **S8**
Figure S5: $^1$H NMR spectrum of tambjamine BE-18591 isolated from *Streptomyces* sp. GSL17-113 **S15**
Figure S6. UV profile and HR-ESI-MS of tambjamine BE-18591 **S16**

**Supplemental References** **S18**

**Table S1.** DADA2 output and taxonomic classification of amplicon sequence variants.
Due to the size of this table, it is available as a separate downloadable CSV file.

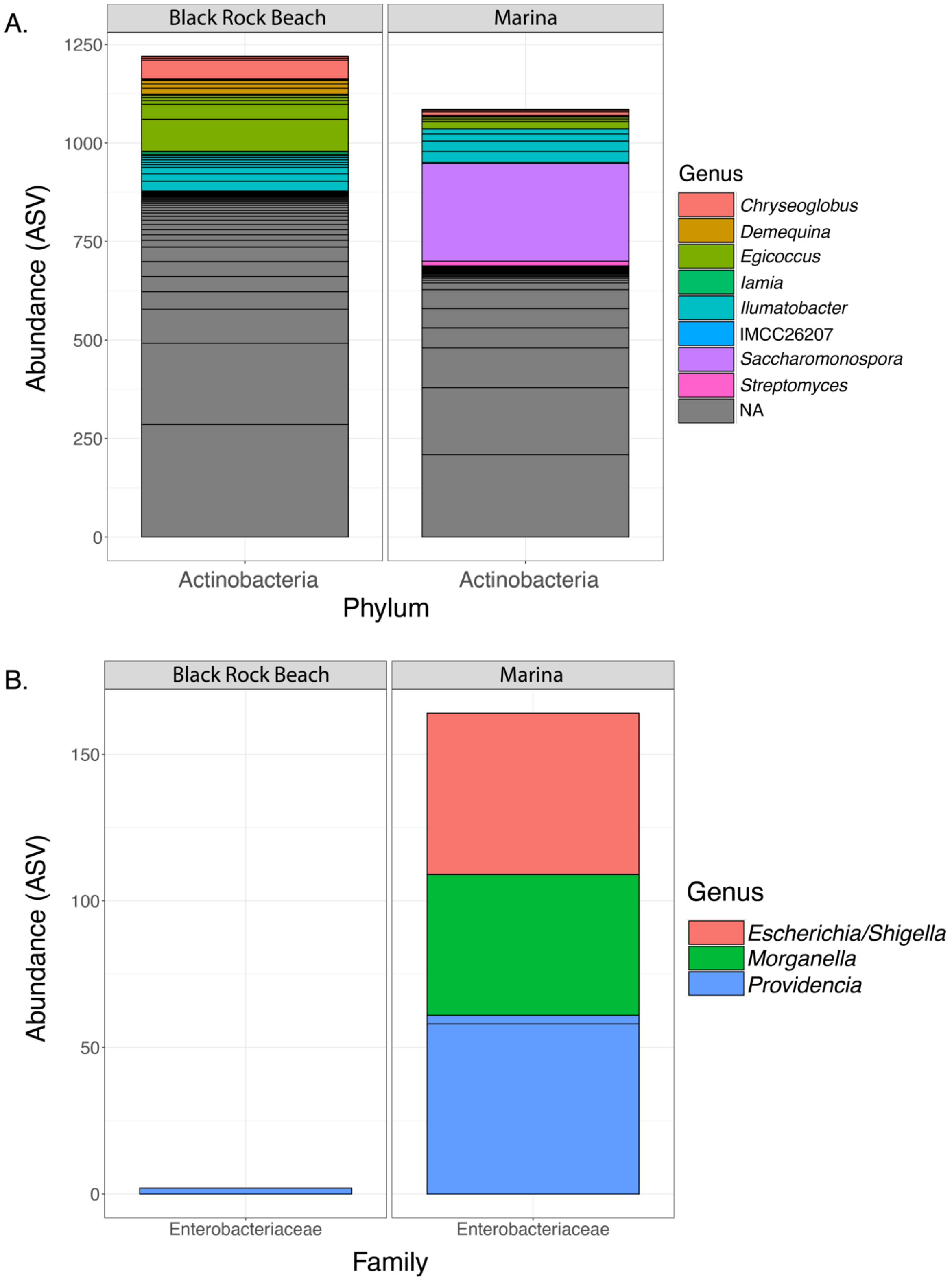


**Figure S1**. Differential abundance of amplicon sequence variants (ASVs) categorized to represent Actinomycetota and Enterobacteriaceae between Black Rock Beach and Marina samples. **A.** Colors represent Actinomycetota genera identified in our analysis. Of these, ASVs categorized to represent *Saccharomonospora* were found at significantly higher abundance in the Marina vs BRB. NA values represent unknown genera. **B.** Differential abundance of Enterobacteriacaea family organisms between BRB and Marina.

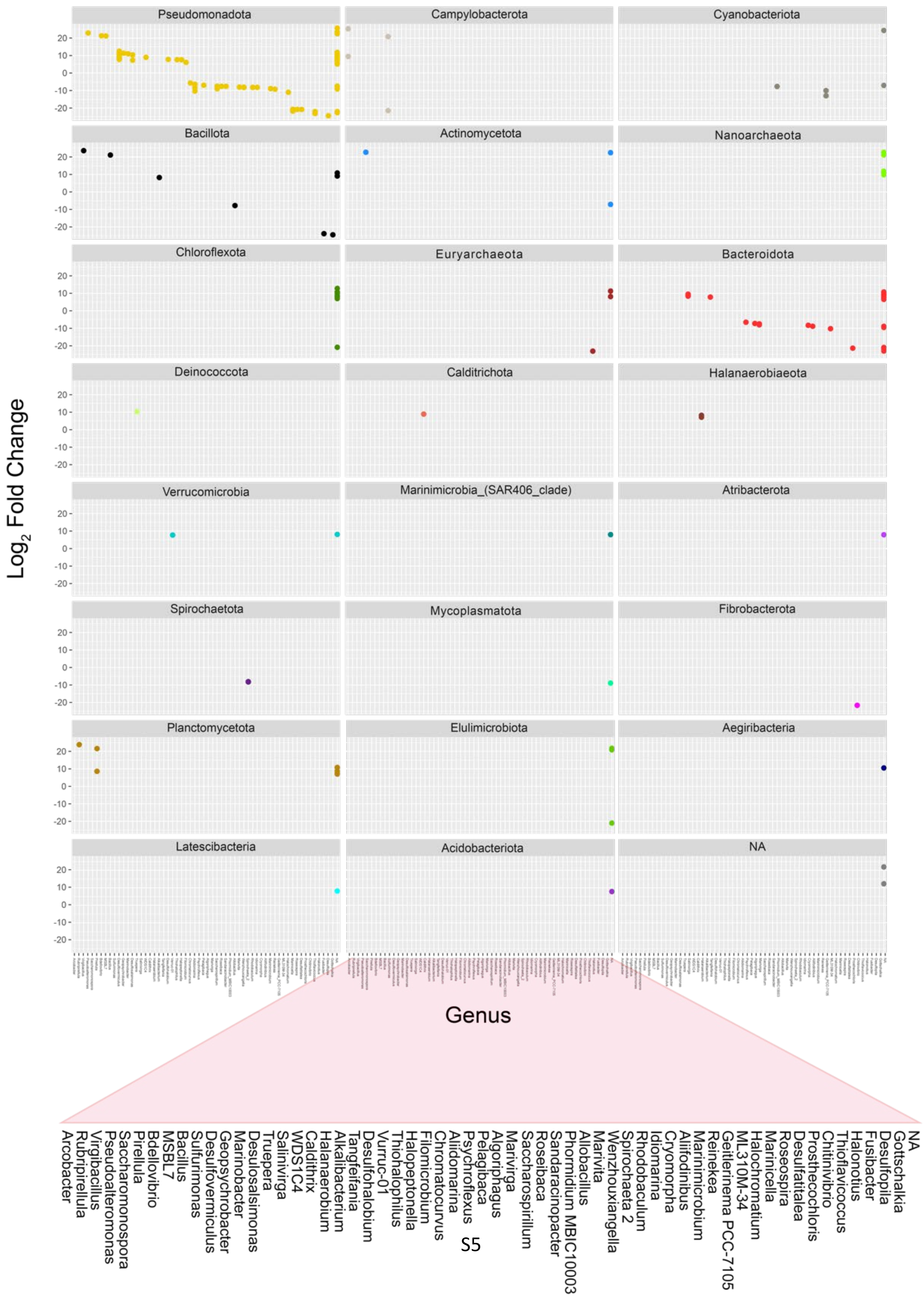
Pseudomonadota
Campylobacterota
Cyanobacteriota
Bacillota
Actinomycetota
Nanoarchaeota
Chloroflexota
Euryarchaeota
Bacteroidota
Deinococcota
Calditrichota
Halanaerobiaeota
Verrucomicrobia
Marinimicrobia_(SAR406_clade)
Atribacterota
Spirochaetota
Mycoplasmatota
Fibrobacterota
Planctomycetota
Elulimicrobiota
Aegiribacteria
Latescibacteria
Acidobacteriota
NA
20
10
0
-10
-20
$Log_2$ Fold Change
Genus
Arcobacter
Rubripirellula
Virgibacillus
Pseudoalteromonas
Saccharomonospora
Pirellula
Bdellovibrio
MSBL7
Bacillus
Sulfurimonas
Desulfovermiculus
Geopsychrobacter
Marinobacter
Desulosalsimonas
Truepera
Salinivirga
WDS1C4
Caldithrix
Halanaerobium
Alkalibacterium
Tangfeifania
Desulfohalobium
Vurruc-01
Thiohalophilus
Halopeptonella
Filomicrobium
Chromatocurvus
Aliidomarina
Psychroflexus
Pelagibaca
Algoriphagus
Marivirga
Saccharospirillum
Roseibaca
Sandaracinopacter
Phormidium MBIC10003
Allobacillus
Marivita
Wenzhouxiangella
Spirochaeta 2
Rhodobaculum
Idiomarina
Cryomorpha
Aliifodinibus
Marinimicrobium
Reinekea
Geitlerinema PCC-7105
ML310M-34
Halochromatium
Marinicella
Roseospira
Desulfatitalea
Prosthecochloris
Chitinivibrio
Thioflavicoccus
Halonotius
Fusibacter
Desulfopila
Gottschalkia
NA

S5

**Figure S2**. Facet wrapped DESeq2 [1] plot showing phyla on the y-axis and genera on the x-axis illustrating differential abundance of microbes between Black Rock Beach and the Marina based on amplicon sequence variants (ASV). Phyla are displayed on the y-axis and genera on the x-axis. A positive log2 fold-change indicates a significantly higher abundance at the Marina, while a negative log2 fold-change indicates significantly higher abundance at Black Rock Beach. Analysis was generated using an alpha value cutoff of < 0.01. NA/NA values represent unknown phylum/unknown genus, and potentially represent uncharacterized microbes. This is an expansion of Figure 3, with each organism assigned its own tile. Each tile is titled by organism phylum. The zoomed in section offers an enlarged view of the x-axis labels, which is the same between all tiles.

**Table S2**. Average nucleotide identity and digital DNA-DNA hybridization analysis of *Streptomyces* sp. GSL17-113 and *Saccharomonospora* sp. GSL17-019. ANI = average nucleotide identity. dDDH = digital DNA-DNA hybridization. C.I. = confidence interval.

| **GSL Strain** | **GSL17-113** | **GSL17-019** |
|---:|---:|---:|
| **Closest Reference Strain (ANI)** | *Streptomyces albus*, subsp. *albus* | [*Actinopolyspora*]_*iraqiensis*_IQ-H1 |
| **Reference Strain Assembly (ANI)** | GCF_000725885 | GCF_000430445 |
| **Estimated ANI** | 0.993 | 0.935 |
| **Reference Strain (dDDH)** | *Streptomyces albus* NBRC 13014 | *Saccharomonospora iraqiensis* subsp. *paurometabolica* YIM90007 |
| **dDDH (d0, in %)** | 90.5 | 42.5 |
| **C.I. (d0, in %)** | [87.3 - 92.9] | 39.1 - 45.9 |
| **dDDH (d4, in %)** | 95.7 | 54.5 |
| **C.I. (d4, in %)** | [94.2 - 96.8] | 51.8 - 57.2 |
| **dDDH (d6, in %)** | 93.7 | 43.9 |
| **C.I. (d6, in %)** | [91.5 - 95.3] | 40.9 - 47.0 |
| **Consensus** | *Streptomyces albus* | N/A |

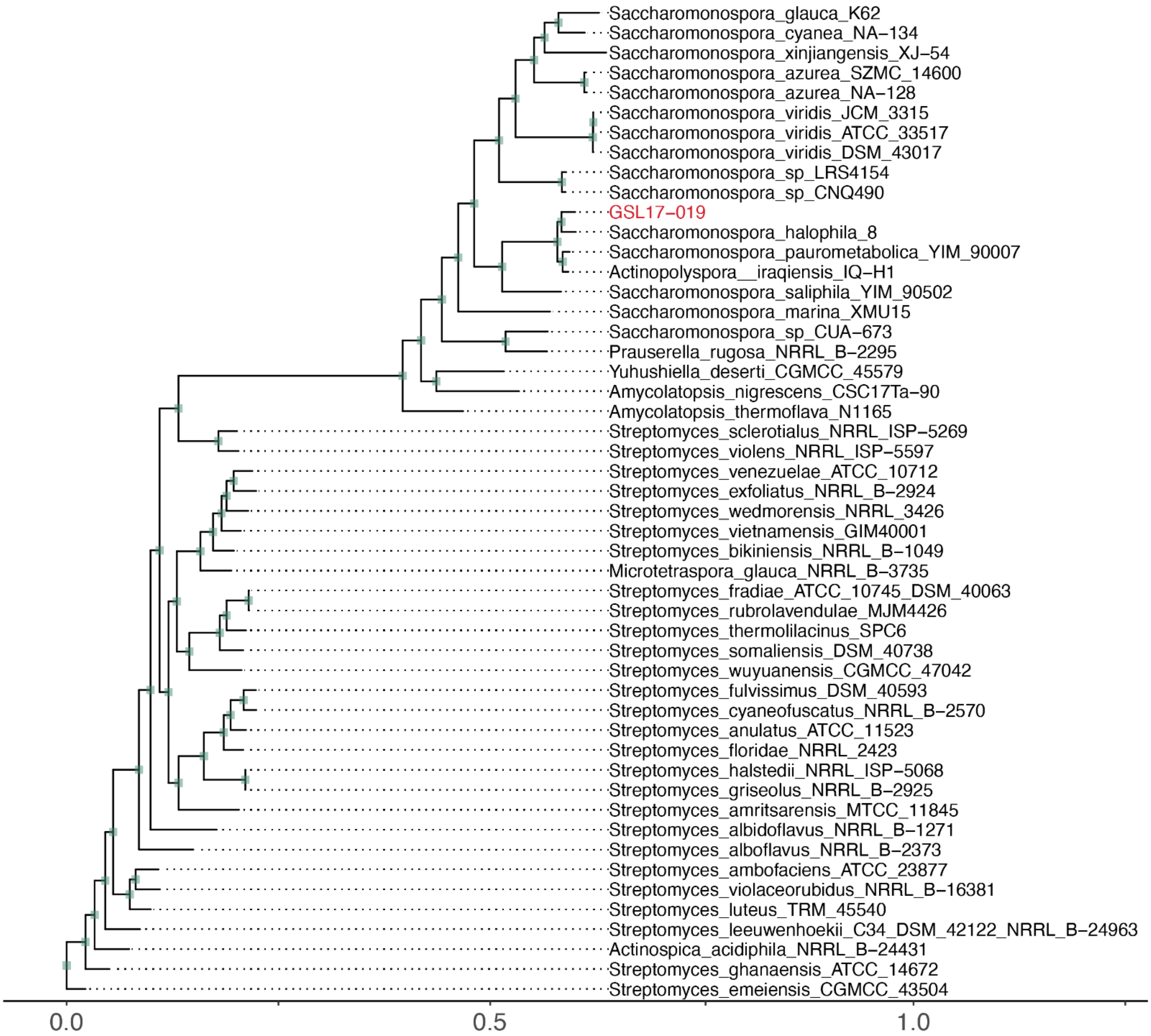


**Figure S3**. Maximum-likelihood phylogenetic tree of *Saccharomonospora* sp. GSL17-019 with reference strains. Phylogenetic analysis was performed using autoMLST with with IQ-TREE [2] Ultrafast Boostrap analysis using 1000 replicates. *Streptomyces emeiensis* CGMCC 43054 was assigned as the outgroup.

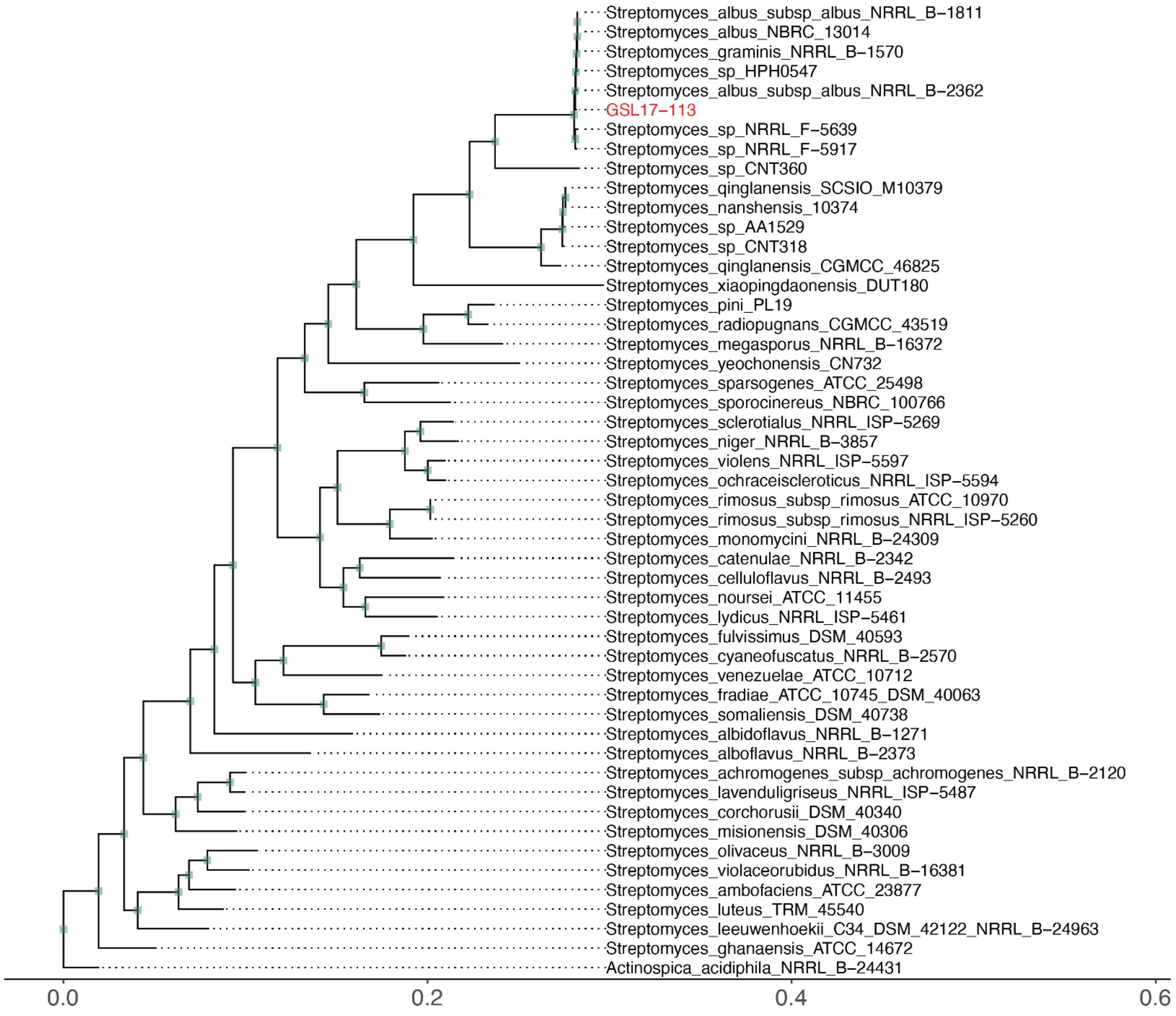


**Figure S4**. Maximum-likelihood phylogenetic tree of *Streptomyces* sp. GSL17-113 with reference strains. Phylogenetic analysis was performed using autoMLST with IQ-TREE [2] Ultrafast Boostrap analysis using 1000 replicates. *Actinocpica acidiphila* NRRL B-24431 was assigned as the outgroup.

**Table S3**. antiSMASH [3] output table of predicted biosynthetic gene clusters identified *Saccharomonospora* sp. GSL17-019.

| Region | Type | From | To | Most similar known cluster | Similarity |
|---|---|---|---|---|---|
| Region 1.1 | indole | 340,734 | 361,840 | fortimicin | 9% |
| Region 1.2 | ectoine | 686,756 | 697,157 | ectoine | 100% |
| Region 1.3 | arylpolyene,ladderane,NRPS | 942,573 | 1,012,591 | coprisamides | 86% |
| Region 2.1 | other,NRPS | 16,861 | 67,002 | lidamycin | 15% |
| Region 2.2 | ranthipeptide | 154,725 | 176,239 | N/A | N/A |
| Region 2.3 | T1PKS | 714,046 | 785,157 | kendomycin B | 15% |
| Region 2.4 | T1PKS,terpene | 850,069 | 909,489 | maduropeptin | 34% |
| Region 3.1 | lanthipeptide-class-i | 132,991 | 158,051 | N/A | N/A |
| Region 3.2 | NRP-metallophore,NRPS | 227,585 | 286,350 | mirubactin | 71% |
| Region 3.3 | terpene | 323,793 | 349,995 | hopene | 46% |
| Region 3.4 | NRPS,T1PKS | 518,191 | 576,596 | caerulomycin A | 8% |
| Region 3.5 | NRPS-like | 688,845 | 729,416 | youssoufenes | 14% |
| Region 3.6 | T3PKS | 753,301 | 794,356 | loseolamycins | 16% |
| Region 3.7 | NRP-metallophore,NRPS | 875,274 | 932,556 | polyoxypeptin | 32% |
| Region 5.1 | terpene | 148,627 | 169,832 | isorenieratene | 36% |
| Region 7.1 | nucleoside | 35,777 | 56,157 | tubercidin | 63% |
| Region 8.1 | other | 33,407 | 64,252 | polyoxypeptin | 16% |
| Region 31.1 | RRE-containing | 1 | 1,629 | N/A | N/A |
| Region 147.1 | RiPP-like | 1 | 1,133 | N/A | N/A |
| Region 193.1 | NRPS | 1 | 1,073 | xenematide | 100% |

**Table S4**. antiSMASH [3] output table of predicted biosynthetic gene clusters identified in *Streptomyces* sp. GSL17-113.

| Region | Type | From | To | Most similar known cluster | Similarity |
|---|---|---|---|---|---|
| Region 2.1 | NRPS-independent-siderophore | 1 | 8,961 | desferrioxamine E | 100% |
| Region 2.2 | terpene | 168,897 | 205,590 | N/A | N/A |
| Region 2.3 | prodigiosin | 463,346 | 498,329 | tambjamine BE-18591 | 96% |
| Region 3.1 | NRPS-independent-siderophore | 54,094 | 69,396 | N/A | N/A |
| Region 3.2 | other,NRPS-like,T1PKS | 211,159 | 274,263 | sanglifehrin A | 18% |
| Region 5.1 | lanthipeptide-class-i | 15,694 | 40,339 | N/A | N/A |
| Region 5.2 | CDPS | 86,778 | 107,614 | a201a | 8% |
| Region 5.3 | other,nucleoside | 301,350 | 342,588 | pseudouridimycin | 68% |
| Region 5.4 | lanthipeptide-class-iii | 359,213 | 381,927 | SapB | 75% |
| Region 7.1 | lassopeptide | 169,240 | 192,222 | aborycin | 64% |
| Region 8.1 | terpene | 191,040 | 213,271 | geosmin | 100% |
| Region 12.1 | T2PKS | 1 | 38,642 | xantholipin | 55% |
| Region 12.2 | ectoine | 194,267 | 204,671 | ectoine | 100% |
| Region 13.1 | T1PKS | 1 | 35,210 | ibomycin | 29% |
| Region 15.1 | NRPS-like | 659 | 25,359 | meilingmycin | 6% |
| Region 15.2 | NRP-metallophore,NRPS | 38,409 | 86,998 | griseobactin | 53% |
| Region 17.1 | NRPS | 1 | 24,264 | dudomycin A | 47% |
| Region 18.1 | T1PKS,oligosaccharide | 1 | 72,435 | ibomycin | 58% |
| Region 21.1 | terpene | 1 | 24,664 | hopene | 61% |
| Region 23.1 | lanthipeptide-class-i, T1PKS,RiPP-like | 10,596 | 62,743 | 4-hexadecanoyl-3-hydroxy-2-(hydroxymethyl)-2H-furan-5-one | 54% |
| Region 25.1 | T1PKS | 4,760 | 55,801 | abyssomicins | 9% |
| Region 27.1 | T1PKS | 1 | 38,903 | reedsmycins | 20% |
| Region 29.1 | T1PKS | 4,666 | 27,498 | ulleungmycin | 5% |
| Region 31.1 | T1PKS | 1 | 23,355 | gargantulides | 26% |
| Region 38.1 | T1PKS | 1 | 3,734 | N/A | N/A |
| Region 42.1 | T1PKS | 1 | 2,991 | N/A | N/A |
| Region 43.1 | T1PKS | 1 | 2,320 | N/A | N/A |

**Table S5**. antiSMASH [3] output table of predicted biosynthetic gene clusters identified in GSL population genome GSL17-111M.

| Region | Type | From | To | Most similar known cluster | Similarity |
|---|---|---|---|---|---|
| Region 2.1 | NRPS-like,terpene | 54,851 | 98,002 | isorenieratene | 87% |
| Region 2.2 | terpene | 165,870 | 188,019 | hopene | 30% |
| Region 2.3 | T1PKS,NRPS-like,NRPS | 405,592 | 489,334 | enduracidin | 18% |
| Region 3.1 | siderophore | 236,346 | 250,973 | ficellomycin | 3% |
| Region 4.1 | terpene | 38,729 | 60,948 | geosmin | 100% |
| Region 5.1 | T1PKS | 1 | 47,195 | lorneic acid A | 23% |
| Region 5.2 | other | 91,654 | 132,754 | N/A | N/A |
| Region 5.3 | CDPS | 288,781 | 309,521 | primycin | 5% |
| Region 6.1 | lanthipeptide-class-iii | 190,135 | 212,807 | SapB | 75% |
| Region 7.1 | T1PKS,thiopeptide,LAP,lanthipeptide-class-iii | 1 | 66,870 | stambomycins | 40% |
| Region 9.1 | T1PKS,PKS-like | 1 | 40,725 | arsono-polyketide | 66% |
| Region 9.2 | ladderane,arylpolyene | 193,464 | 232,547 | atratumycin | 31% |
| Region 10.1 | NRPS-like | 12,669 | 56,523 | indigoidine | 27% |
| Region 11.1 | transAT-PKS,PKS-like | 1 | 67,362 | lagriamide | 9% |
| Region 11.2 | NRPS,T1PKS,ectoine | 82,641 | 147,540 | ectoine | 100% |
| Region 17.1 | T1PKS,NRPS | 1 | 129,375 | stambomycins | 60% |
| Region 18.1 | siderophore | 93,557 | 109,805 | macrotetrolide | 33% |
| Region 19.1 | lanthipeptide-class-iv | 90,862 | 113,609 | a201a | 5% |
| Region 21.1 | lassopeptide | 1 | 22,102 | anantin C | 75% |
| Region 21.2 | NRPS-like | 60,014 | 102,755 | meridamycin | 13% |
| Region 22.1 | NRPS-like,lanthipeptide-class-i | 1,396 | 44,440 | saquayamycin A | 7% |
| Region 22.2 | NRPS | 85,478 | 113,360 | diisonitrile antibiotic SF2768 | 66% |
| Region 24.1 | ectoine | 81,876 | 92,262 | N/A | N/A |
| Region 25.1 | T1PKS | 4,705 | 57,717 | pyrrolomycins | 42% |
| Region 26.1 | NRPS,lanthipeptide-class-i,lanthipeptide-class-ii | 16,299 | 70,052 | meilingmycin | 3% |
| Region 27.1 | CDPS | 42,666 | 63,502 | daptomycin | 6% |
| Region 29.1 | lanthipeptide-class-i | 31,831 | 56,281 | N/A | N/A |
| Region 30.1 | RiPP-like | 8,308 | 20,209 | N/A | N/A |
| Region 39.1 | T2PKS,RRE-containing,T1PKS,lanthipeptide-class-i | 1 | 57,225 | xantholipin | 18% |
| Region 42.1 | T1PKS | 1 | 31,923 | 5-acetyl-5,10-dihydrophenazine-1-carboxylic acid / 5-(2-hydroxyacetyl)-5,10-dihydrophenazine-1-carboxylic acid / endophenazines | 13% |

| | | | | | |
|---|---|---|---|---|---|
| Region 45.1 | NRPS | 26,174 | 54,793 | stenothricin | 9% |
| Region 49.1 | thiopeptide,LAP | 9,680 | 39,632 | kistamicin A | 16% |
| Region 51.1 | NRPS,NRPS-like | 1 | 37,944 | himastatin | 12% |
| Region 56.1 | NRPS-like,NRPS | 1 | 35,917 | limazepines | 55% |
| Region 57.1 | T1PKS | 13,687 | 45,652 | herbimycin A | 6% |
| Region 58.1 | NRPS | 1 | 41,724 | N/A | N/A |
| Region 59.1 | lanthipeptide-class-i,lassopeptide | 1 | 39,784 | aborycin | 57% |
| Region 62.1 | RiPP-like | 35,126 | 43,610 | N/A | N/A |
| Region 64.1 | T1PKS,NRPS-like,prodigiosin | 1 | 42,884 | marineosins | 100% |
| Region 70.1 | T1PKS | 1 | 29,398 | N/A | N/A |
| Region 78.1 | T1PKS | 1 | 28,673 | 4-hexadecanoyl-3-hydroxy-2-(hydroxymethyl)-2H-furan-5-one | 45% |
| Region 79.1 | thiopeptide,LAP | 8,998 | 37,084 | nocathiacin | 4% |
| Region 81.1 | terpene | 3,184 | 25,466 | hygromycin A | 6% |
| Region 85.1 | NRPS,T1PKS | 153 | 36,407 | N/A | N/A |
| Region 86.1 | lanthipeptide-class-i | 1 | 20,633 | N/A | N/A |
| Region 95.1 | terpene | 1 | 20,406 | isorenieratene | 85% |
| Region 126.1 | T1PKS | 1 | 29,207 | sceliphrolactam | 24% |
| Region 133.1 | T1PKS | 1 | 27,868 | nigericin | 88% |
| Region 134.1 | other,nucleoside | 1 | 24,408 | pseudouridimycin | 68% |
| Region 145.1 | terpene | 1 | 17,302 | N/A | N/A |
| Region 158.1 | T1PKS | 1 | 24,670 | polyoxypeptin | 8% |
| Region 167.1 | T1PKS | 1 | 23,167 | ebelactone | 13% |
| Region 183.1 | T3PKS | 1 | 21,583 | flaviolin | 75% |
| Region 192.1 | terpene | 739 | 20,703 | geosmin | 100% |
| Region 222.1 | T1PKS | 1 | 18,966 | primycin | 18% |
| Region 232.1 | terpene | 3,636 | 18,622 | siomycin A | 7% |
| Region 233.1 | RiPP-like | 10,578 | 18,560 | N/A | N/A |
| Region 246.1 | RRE-containing | 1 | 12,365 | aclacinomycin | 15% |
| Region 273.1 | T1PKS | 1 | 16,190 | butyrolactol A | 26% |
| Region 279.1 | T1PKS,NRPS-like,other | 1 | 15,829 | catenulisporolides | 7% |
| Region 302.1 | RRE-containing | 1 | 15,034 | N/A | N/A |
| Region 314.1 | RRE-containing | 1 | 14,699 | granaticin | 8% |
| Region 389.1 | lanthipeptide-class-iii | 1 | 12,457 | SapB | 75% |
| Region 391.1 | terpene | 1 | 12,441 | N/A | N/A |
| Region 392.1 | siderophore | 1 | 12,426 | N/A | N/A |
| Region 401.1 | NRPS-like | 1 | 12,222 | 2'-chloropentostatin / 2'-amino-2'-deoxyadenosine | 6% |
| Region 413.1 | siderophore | 388 | 11,972 | ficellomycin | 3% |
| Region 423.1 | T1PKS | 1 | 11,809 | niphimycins C-E | 29% |
| Region 428.1 | T1PKS | 1 | 11,754 | piericidin A1 | 91% |
| Region 431.1 | T1PKS | 1 | 11,724 | candicidin | 28% |

| | | | | | |
|---|---|---|---|---|---|
| Region 456.1 | terpene | 1 | 10,955 | hopene | 46% |
| Region 459.1 | T1PKS | 1 | 10,889 | salinomycin | 18% |
| Region 555.1 | T1PKS | 1 | 9,019 | heronamides | 25% |
| Region 561.1 | T1PKS | 1 | 8,918 | filipin | 38% |
| Region 600.1 | oligosaccharide | 1 | 8,214 | ibomycin | 15% |
| Region 606.1 | T1PKS | 1 | 8,157 | ibomycin | 7% |
| Region 617.1 | T1PKS | 1 | 7,963 | polyoxypeptin | 10% |
| Region 675.1 | siderophore | 1 | 7,258 | desferrioxamine E | 100% |
| Region 686.1 | RRE-containing | 1 | 7,206 | SSV-2083 | 36% |
| Region 690.1 | T1PKS | 1 | 7,139 | N/A | N/A |
| Region 692.1 | T1PKS | 1 | 7,129 | nanchangmycin | 30% |
| Region 777.1 | terpene | 1 | 6,036 | hopene | 30% |
| Region 801.1 | T1PKS | 1 | 5,827 | N/A | N/A |
| Region 803.1 | T1PKS | 1 | 5,814 | maklamicin | 6% |
| Region 822.1 | T1PKS | 1 | 5,601 | mediomycin A | 28% |
| Region 891.1 | T1PKS | 1 | 4,944 | polyoxypeptin | 10% |
| Region 988.1 | T1PKS | 1 | 4,199 | N/A | N/A |
| Region 1015.1 | terpene | 1 | 4,010 | N/A | N/A |
| Region 1016.1 | T1PKS | 1 | 4,006 | N/A | N/A |
| Region 1066.1 | T1PKS | 1 | 3,651 | N/A | N/A |
| Region 1074.1 | T1PKS | 1 | 3,617 | N/A | N/A |
| Region 1119.1 | NRPS | 1 | 3,382 | N/A | N/A |
| Region 1143.1 | terpene | 1 | 3,258 | hopene | 23% |
| Region 1167.1 | T1PKS | 1 | 3,115 | N/A | N/A |
| Region 1188.1 | siderophore | 1 | 3,053 | desferrioxamine E | 75% |
| Region 1196.1 | terpene | 1 | 3,017 | N/A | N/A |
| Region 1230.1 | T1PKS | 1 | 2,810 | N/A | N/A |
| Region 1335.1 | T2PKS | 1 | 2,349 | xantholipin | 6% |
| Region 1406.1 | T1PKS | 1 | 2,011 | N/A | N/A |
| Region 1505.1 | T1PKS | 1 | 1,634 | N/A | N/A |
| Region 1673.1 | T1PKS | 1 | 1,308 | N/A | N/A |
| Region 1676.1 | T1PKS | 1 | 1,302 | N/A | N/A |
| Region 1749.1 | ectoine | 1 | 1,192 | ectoine | 50% |
| Region 1828.1 | terpene | 1 | 1,071 | N/A | N/A |

**Table S6.** $^{1}$H-NMR resonances of tambjamine BE-18591, isolated from *Streptomyces* sp. GSL17-113. $^{1}$H chemical shifts are referenced to CDCL₃.

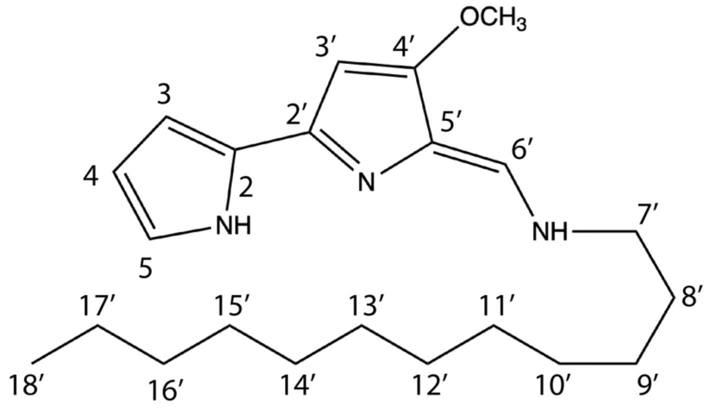


tambjamine BE-18591 500 MHz, $CDCl_3$

| Position | $\delta$ ($^{1}$H), mult (*J* in Hz) |
|---|---|
| 2 | – |
| 3 | 6.74, m |
| 4 | 6.27, m |
| 5 | 7.09, s |
| 2′ | – |
| 3′ | 5.98, s |
| -$OCH_3$ | 3.92, s |
| 4′ | – |
| 5′ | – |
| 6′ | 7.32, d (14.8) |
| 7′ | 3.45, t (6.54) |
| 8′ | 1.70, m |
| 9′ | 1.35, m |
| 10′ | 1.24, o.l.[a] |
| 11′ | 1.24, o.l.[a] |
| 12′ | 1.24, o.l.[a] |
| 13′ | 1.24, o.l.[a] |
| 14′ | 1.24, o.l.[a] |
| 15′ | 1.24, o.l.[a] |
| 16′ | 1.24, o.l.[a] |
| 17′ | 1.24, o.l.[a] |
| 18′ | 0.86, t (6.94) |

[a]o.l. = overlapping signal

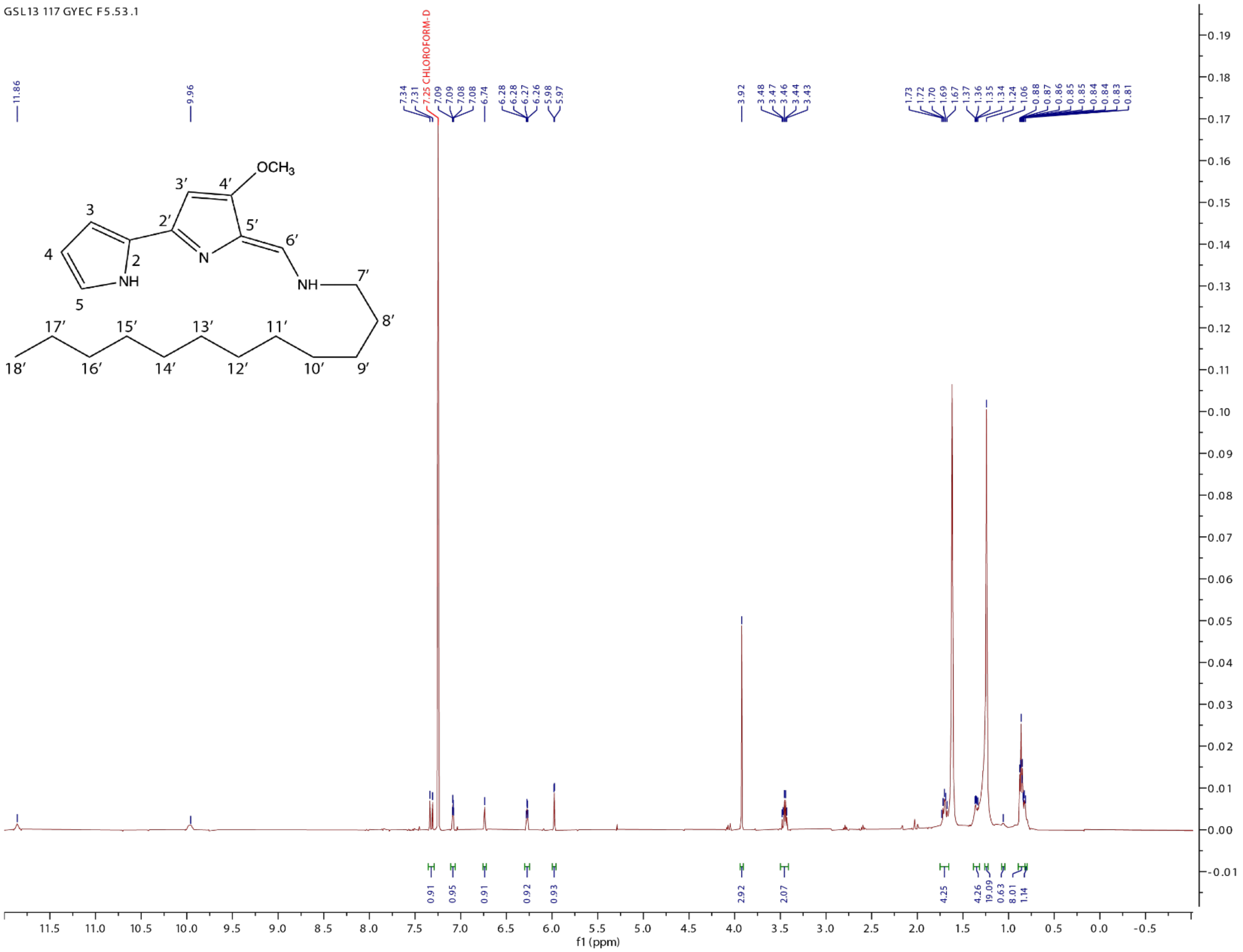

**Figure S5**. $^{1}H$ NMR spectra of tambjamine BE-18591 in $CDCl_3$ isolated from *Streptomyces* sp. GSL17-113.

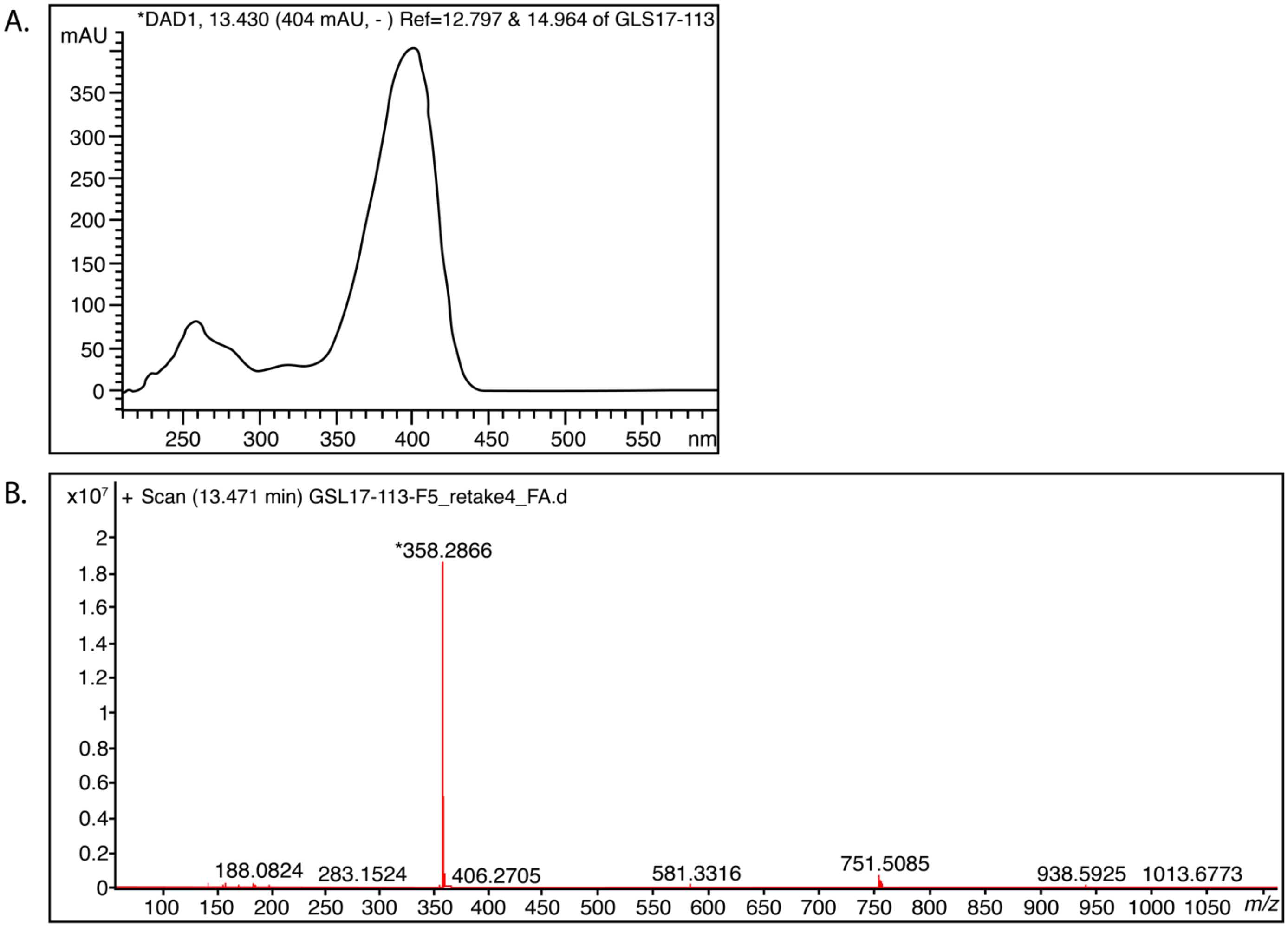


**Figure S6.** Identification of tambjamine BE-18591 from *Streptomyces* sp. GSL17-113. **A)** UV profile and **B)** HR-ESI-MS of tambjamine BE-18591. $C_{22}H_{35}N_3O$ [HRMS *m/z*: $[M+H]^+$ calcd for $C_{22}H_{36}N_3O$ 358.2858; found 358.2866.

**Table S7**. Putative functions of the gene products of the tambjamine BE-18591 biosynthetic gene cluster in *Streptomyces* sp. GSL17-113.

| Gene Product | Total Amino Acids | Proposed Function | Sequence Similarity (Origin) | Similarity/ Identity (%) | Accession Number |
|---|---|---|---|---|---|
| TabA | 448 | Aminotransferase class-III | *Streptomyces sp.* NRRL F-5639 | 95/99.5 | WP_031025583.1 |
| TabC | 162 | SRPBCC family protein | *Streptomyces* | 99/100 | WP_016467651.1 |
| TabD | 254 | AfsR/SARP family transcriptional regulator | *Streptomyces* | 99/99.6 | WP_016467654.1 |
| TabE | 611 | Aldehyde dehydrogenase | *Streptomyces albus* | 99/99.5 | TGG88512.1 |
| TabH | 886 | PEP/pyruvate-binding domain-containing protein | *Streptomyces* | 99/100 | WP_016467639.1 |
| TabI | 355 | Class I SAM-dependent methyltransferase | *Streptomyces* | 99/100 | WP_016467640.1 |
| TabJ | 280 | Alpha/beta fold hydrolase | *Streptomyces* | 99/99.6 | WP_016467641.1 |
| TabM | 538 | D-alanine--poly(phosphoribitol) ligase | *Streptomyces* | 99/100 | WP_016467642.1 |
| TabN | 630 | Aminotransferase class I/II-fold pyridoxal phosphate-dependent enzyme | *Streptomyces* | 99/99.8 | WP_031174881.1 |
| TabO | 86 | Acyl-carrier protein | *Streptomyces* | 98/100 | WP_016467644.1 |
| TabP | 337 | β-ketoacyl-ACP synthase | *Streptomyces* | 99/100 | WP_016467650.1 |
| TabQ | 78 | Acyl-carrier protein | *Streptomyces* | 98/100 | WP_016467649.1 |
| TabR | 413 | β-ketoacyl-ACP synthase | *Streptomyces sp.* NRRL F-5639 | 99/99.5 | WP_031025584.1 |
| TabU | 243 | 4'-phosphopantetheinyl transferase superfamily protein | *Streptomyces* | 86/100 | WP_041968511.1 |
| TabV | 414 | Oxidase/dehydrogenase | *Streptomyces* sp. NRRL F-5917 | 94/100 | WP_107047698.1 |
| TabW | 380 | Acyl-CoA dehydrogenase | *Streptomyces* | 99/100 | WP_016467646.1 |
| TabX | 986 | Polyketide Synthase (KS-KS) | *Streptomyces albus* | 99/99.9 | WP_037611838.1 |
| TabY | 105 | Hypothetical | *Streptomyces* | 99/100 | WP_016467647.1 |

**Supplemental References**

1. Love MI, Huber W, Anders S. Moderated estimation of fold change and dispersion for RNA-seq data with DESeq2. *Genome Biol*. 2014; 15(12):550.

2. Alanjary M, Steinke K, Ziemert N. AutoMLST: an automated web server for generating multi-locus species trees highlighting natural product potential. *Nucleic Acids Res*. 2019; 47(W1):W276-W82.

3. Blin K, Shaw S, Steinke K, Villebro R, Ziemert N, Lee SY, et al. antiSMASH 5.0: updates to the secondary metabolite genome mining pipeline. *Nucleic Acids Res*. 2019; 47(W1):W81-W7.
